# Extracellular vesicles induce aggressive lung cancer via non-canonical integrin-EGFR-KRAS signaling

**DOI:** 10.1101/2022.08.16.504078

**Authors:** Karla Rubio, Addi J. Romero-Olmedo, Pouya Sarvari, Stefan Günther, Aditi Mehta, Birgit Bassaly, Peter Braubach, Gergana Dobreva, Malgorzata Wygrecka, Stefan Gattenlöhner, Thomas Braun, Achim Tresch, Johannes Graumann, Guillermo Barreto

## Abstract

Small cell lung cancer (SCLC) is an extremely aggressive lung cancer type with a patient median survival of 6-12 months. Epidermal growth factor (EGF) signaling plays an important role in triggering SCLC. In addition, growth factor-dependent signals and alpha-, beta-integrin (ITGA, ITGB) heterodimer receptors functionally cooperate and integrate their signaling pathways. However, the precise role of integrins in EGF receptor (EGFR) activation in SCLC has remained elusive. We analyzed RNA-sequencing data, human precision-cut lung slices (hPCLS), retrospectively collected human lung tissue samples and cell lines to demonstrate that non-canonical ITGB2 signaling activates EGFR and RAS/MAPK/ERK signaling in SCLC. Further, we identified a novel SCLC gene expression signature consisting of 93 transcripts that were induced by ITGB2, which might be used for stratification of SCLC patients, prognosis prediction of LC patients and development of patient-tailored therapies. We also found by proteomic analysis a cell-cell communication mechanism based on extracellular vesicles (EVs) containing ITGB2, which were secreted by SCLC cells and induced in control human lung tissue RAS/MAPK/ERK signaling and SCLC markers. We uncovered a mechanism of ITGB2-mediated EGFR activation in SCLC that explains EGFR-inhibitor resistance independently of EGFR mutations, suggesting the development of therapies targeting ITGB2 for patients with this extremely aggressive lung cancer type.

## Introduction

Lung cancer (LC) causes more deaths worldwide than the next three most prevalent cancers together (colorectal, breast and prostate) ^1^. Based on histology, LC is classified into non-small cell (NSCLC) and small cell lung cancer (SCLC). NSCLC can be further classified into three subtypes: squamous cell carcinoma, adenocarcinoma, and large-cell lung cancer ^2^. SCLC accounts for approximately 15-20% of all LC cases and is strongly associated with cigarette smoking ^3^. SCLC is a neuroendocrine type of lung cancer characterized by aggressive progression due to high cellular proliferation and early metastasis ^4^. The first line of therapy includes a combination of platinum-based treatment (cisplatin or carboplatin) with etoposide or irinotecan. While SCLC is initially chemo- and radiosensitive, therapy resistance frequently arises. Consequently, patient prognosis is poor with a median survival of 6 to 12 months ^5^. Extracellular vesicles (EVs) are nano-sized, phospholipid membrane-enclosed vesicles that are secreted by different cell types into the extracellular space and can be found in a wide spectrum of human body fluids including serum, plasma, saliva, breast milk, amniotic fluid, cerebrospinal fluid, urine, among others ^6, 7^. EVs are classified based on their size, biogenesis and method of cellular release into three groups: exosomes, microvesicles and apoptotic bodies. Microvesicles and apoptotic bodies are formed by budding from the plasma membrane, and generally range in size from 0.1 to 1 μm and 1 to 4 μm, respectively ^8, 9^. In contrast, exosomes are smaller with a diameter ranging from 30 to 150 nm, and are formed by inward budding of the endosome lumen to form a multivesicular body, which fuses with the plasma membrane during secretion ^10^. Due to an overlap in size (100–150 nm) and density (1.08–1.19 g/ml), as well as the presence of common markers, such as CD63 and CD81 ^11^, it is frequently difficult to differentiate exosomes and microvesicles. Thus, exosomes and microvesicles with a diameter below 150 nm are collectively referred to as small extracellular vesicles (small EVs)^12^. EV secretion is elevated in response to inflammation ^13^ and hypoxia ^14, 15^ and is associated with human diseases. For example in cancer, EV levels correlate with tumor invasiveness ^16–18^. Interestingly, EVs contain proteins and nucleic acids, and it has been reported that tumor cells can influence their microenvironment through EV-based cell-cell communication mechanisms^19–23^.

Epidermal growth factor (EGF) signaling plays an important role in LC development and metastasis. Particularly, activation of EGF receptor (EGFR) tyrosine kinases (RTK) is crucial for triggering SCLC and NSCLC ^24^. EGFR is composed of an extracellular ligand-binding domain, a transmembrane domain and an intracellular tyrosine kinase domain. The binding of a ligand to the extracellular domain of EGFR induces receptor dimerization, activation of the intracellular kinase domain and auto-phosphorylation of tyrosine residues within the cytoplasmic domain of the receptor ^25^. The tyrosine-phosphorylated motifs of EGFR recruit adapters or signaling molecules that initiate various downstream signaling cascades including the RAS/MAPK/ERK, PIK3-Akt and STAT pathways ^26^. These signaling cascades result in transcriptional activation of gene signatures that mediate specific cellular responses, such as cell proliferation, cell migration, epithelial-mesenchymal transition (EMT), among others.

Integrins are heterodimeric transmembrane protein complexes resulting from noncovalent association of specific alpha (ITGA) and a beta (ITGB) subunits ^27^. In general, each integrin subunit has a large extracellular domain, a single-pass transmembrane domain, and a short cytoplasmic tail ^27^. Integrins mediate cell−cell and cell−ECM (extracellular matrix) interactions and transmit signals in both directions, outside-in and inside-out across the cell membrane ^28^. Recent studies have shown that growth factor- and integrin-dependent signals functionally cooperate to integrate their signaling pathways ^29, 30^. The crosstalk between EGFR and integrins has been reported to play a key role in multiple biological processes in cancer ^31^. Moreover, several integrin dimers, including ITGA5-ITGB1, ITGA6-ITGB4, ITGAv-ITGB3 and ITGAv-ITGB5, exert different effects on the regulation of EGF signaling ^32–34^. We have previously demonstrated that ITGA2, ITGB2 and ITGB6 are enriched in the membrane of alveolar type-II (ATII) cells, which are lung progenitor cells responsible for regeneration of the alveolar epithelium during homeostatic turnover and in response to injury ^35^. In addition, ATII cells were also reported as cells of origin of lung adenocarcinoma ^36^. These observations motivated us to investigate the function of integrin receptor subunits in LC.

## Results

### Non-canonical ITGB2 signaling activates EGFR in SCLC

The previously reported enrichment of ITGA2, ITGB2 and ITGB6 in the membrane of lung progenitor cells ^35^ suggests a potential interaction between these integrin receptor subunits. To test this hypothesis, we performed *in silico* analyses of proteomic data repositories derived from protein-protein interaction databases (Figure S1A). Using the STRING database, we identified 50 interaction partners of human ITGA2 with high confidence (combined score ≥ 0.9; 2 nodes, Table S1), including ITGB2 (combined score=0.96) and ITGB6 (combined score=0.97). To confirm the interaction of ITGA2 with ITGB2 and ITGB6, we performed co-immunoprecipitation (Co-IP) assays using total protein extracts from mouse lung epithelial cells (MLE-12) transiently transfected with *HIS-ITGA2* and *YFP*-*ITGB2* or *GFP-ITGB6* (Figure 1A). ITGA2 precipitated both ITGB2 and ITGB6, thereby validating our *in silico* analysis and demonstrating the interaction between these integrin receptor subunits. To assess whether *ITGB2*, *ITGB6* and *ITGA2* are expressed in SCLC, we performed qRT-PCR-based expression analysis using total RNA extracted from retrospectively collected formalin-fixed paraffin embedded (FFPE) human lung tissues from SCLC patients (*n*=5) and control donors (Ctrl, *n*=4; Table S2 and Figure 1B). We observed increased *ITGB2* (*P*=0.02) and *ITGA2* (*P*=9E-3) expression in SCLC as compared to Ctrl FFPE human lung tissue, whereas *ITGB6* levels were reduced (*P*=0.01). To perform an equivalent analysis in NSCLC, we retrieved RNA-sequencing (RNA-seq) data of lung adenocarcinoma patients (LUAD, *n*=11) and control donors (Ctrl, *n*=9) from The Cancer Genome Atlas (TCGA) (Table S3). In contrast to SCLC, we observed decreased *ITGB2* levels (*P*=0.05) in LUAD as compared to Ctrl human lung tissue (Figure 1C). Similarly to SCLC, *ITGA2* expression also increased (*P*=0.01) in LUAD compared to Ctrl. Consistent with these results, we found a positive correlation between the expression of *ITGB2* and *ITGA2* in SCLC (R^2^=0.84, *P*<0.05, Figure S1B, top) and a positive correlation between the expression of *ITGB6* and *ITGA2* in LUAD (R^2^=0.94, *P*<1E-4, Figure S1B, bottom) by linear regression analysis. Our results demonstrated the complementary expression of *ITGB2* and *ITGB6* in SCLC and LUAD suggesting their use as markers for these cancer subtypes and supporting the formation of different integrin heterodimer receptors in SCLC and LUAD.

**Figure 1:**
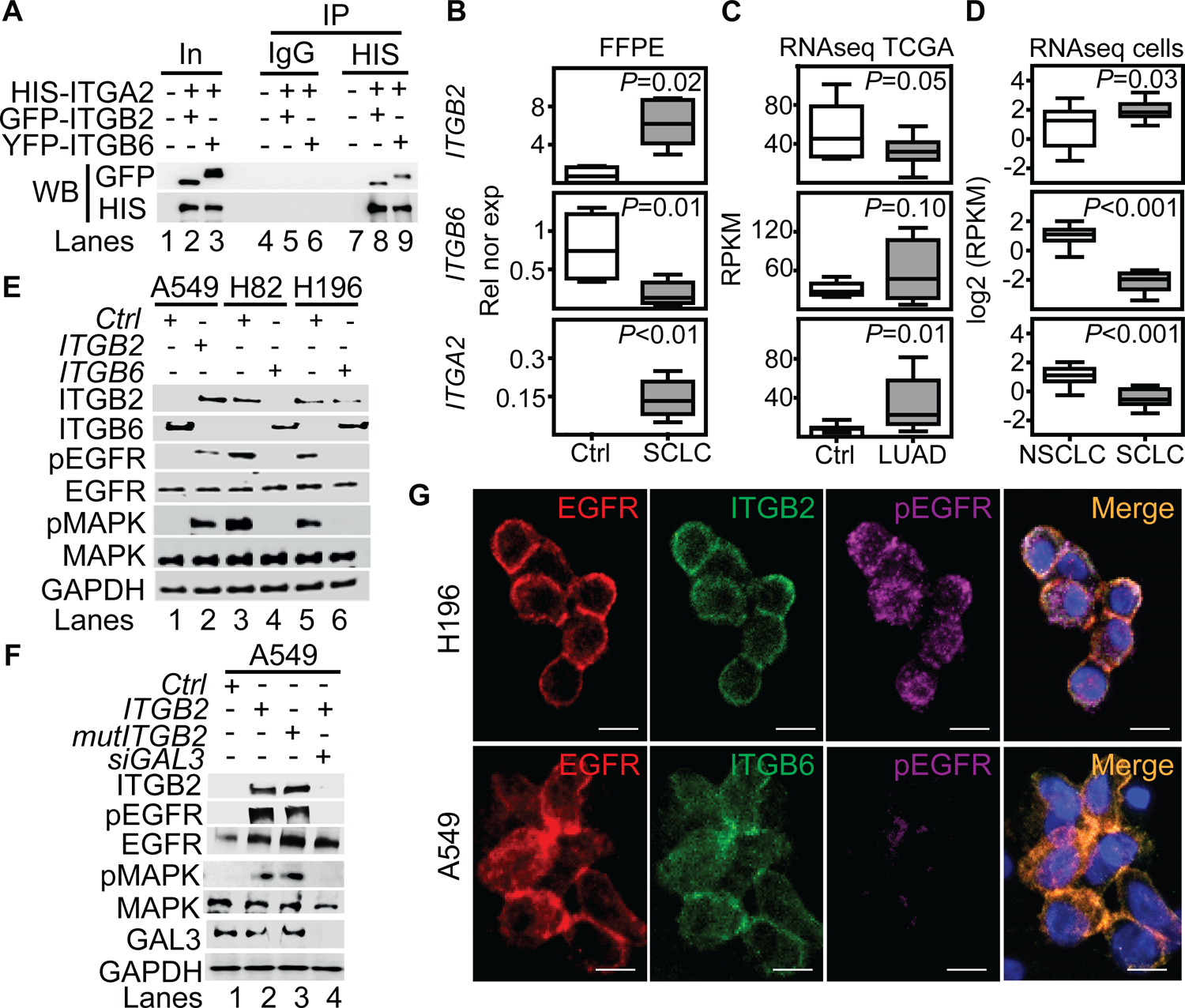
Non-canonical ITGB2 signaling activates EGFR in SCLC. (**A**) Protein extracts of MLE-12 cells co-transfected with *ITGA2*-*HIS* and *ITGB2*-*YFP* or *ITGA2*-*HIS* and *ITGB6*-*GFP* were immunoprecipitated (IP) using either immunoglobulin G (IgG, as control) or HIS-specific antibodies. Co-IP proteins were analyzed by WB using the indicated antibodies. In, input, 3% of material used for the IP. (**B**) Box plots of qRT-PCR-based expression analysis of indicated transcripts using RNA isolated from FFPE lung tissue sections from Ctrl (*n*=4) and small cell-lung cancer (SCLC, *n*=5) patients. Rel nor exp, relative normalized expression to *GAPDH*. (**C**) Box plots of RNA-seq-based expression analysis of indicated transcripts in matched control donors (Ctrl; *n*=9) and matched lung adenocarcinoma (LUAD; *n*=11) patients from the Cancer Genome Atlas (TCGA). Values were normalized using reads per kilobase per million (RPKM). (**D**) Box plots of RNA-seq-based expression analysis of indicated transcripts in non-small cell lung cancer (NSCLC; *n*=33) and small cell lung cancer (SCLC; *n*=17) cell lines. Values are represented as log2 RPKM. All box plots (C-D) indicate median (middle line), 25th, 75th percentile (box) and 5th and 95th percentile (whiskers); *P*-values after two-tailed t-test. Source data are provided as Source Data S1. (**E**) Total protein extracts of A549, NCI-H82 and NCI-H196 cell lines transfected with *ITGB2* or *ITGB6* were analyzed by WB using the indicated antibodies. (**F**) Total protein extracts of A549 cells transfected with *ITGB2,* ligand-binding-deficient D134A ITGB2 mutant (*mutITGB2*) or Galectin-3-specific small interfering RNA (*siGAL3*) were analyzed by WB using the indicated antibodies. (**G**) Confocal microscopy after immunostaining with specific antibodies against EGFR, pEGFR, ITGB2 and ITGB6 in NCI-H196 and A549 cells. DAPI, nucleus. Scale bars, 10 μm. See also Figure S1-S6.

To further investigate the differential expression of *ITGB6* and *ITGB2* in lung cancer subtypes, we analyzed publicly available RNA-seq data of NSCLC and SCLC cell lines (Figure S2) ^37^. Consistent with our results in human lung tissue (Figure 1B-C), we detected high levels of *ITGB2* in SCLC cell lines (Figure 1D, top), whereas *ITGB6* levels were high in NSCLC cell lines (Figure 1D, middle). In addition, linear regression analysis confirmed the positive correlation between *ITGB2* and *ITGA2* levels in SCLC cell lines (R^2^=0.82; *P*=4E-3; Figure S1C, top) and the positive correlation between *ITGB6* and *ITGA2* levels in NSCLC cell lines (R^2^=0.72; *P*=9E-4; FigureS1C, bottom). To further investigate these findings, we selected the human adenocarcinoma cell line A549 as representative of NSCLC cells, whereas the human cell lines NCI-H82 and NCI-H196 were selected as representative cells for SCLC. The selected cell lines lack somatic mutations in the *EGFR* locus (Figure S3) and are experimental systems commonly used in LC subtype-specific studies. Expression analysis by qRT-PCR (Figure S4A-B) confirmed the results obtained by RNA-seq (Figure 1D). Moreover, our qRT-PCR-based expression analyses were also confirmed by Western Blot analysis (WB) of protein extracts from transfected A549, NCI-H82 and NCI-H196 cells (Figure 1E, top), and in sections of FFPE lung tissue from NSCLC and SCLC patients using ITGB6- or ITGB2-specific antibodies (Figure S4C-D). Our results support a mutual negative regulation of *ITGB6* and *ITGB2* expression in NSCLC and SCLC.

Additional analysis of the RNA-seq data from NCI-H82 and NCI-H196 cells showed enrichment of pathways related to EGFR signaling (Figure S5A-B). Further, we observed increased levels of key downstream genes of EGFR signaling, such as *VIM*, *NFKB2* and *HIF1A*, in RNA-seq data from SCLC cell lines when compared to NSCLC cell lines (Figure S5C-D) ^37^, which were confirmed by qRT-PCR-based expression analysis in FFPE human lung tissue (Figure S5E). Our results indicate that EGFR signaling is active in SCLC and correlates with increased *ITGB2* expression. To further investigate this finding, we analyzed protein extracts from transiently transfected A549, NCI-H82 and NCI-H196 cells by WB (Figure 1E, middle). Overexpression of *ITGB2* in A549 cells induced phosphorylation of EGFR (pEGFR) and the mitogen-activated protein kinase (pMAPK) as compared to *Ctrl* transfected cells. On the other hand, the levels of pEGFR and pMAPK in NCI-H82 and NCI-H196 cells were reduced after *ITGB6* transfection. Moreover, the changes in pEGFR and pMAPK in all three cells lines occurred without affecting total levels of EGFR and MAPK, thereby indicating that the observed effects were related to the post-translational phosphorylation of these proteins. To gain further insights into the mechanism of ITGB2-induced, phosphorylation-dependent activation of EGFR and MAPK, we investigated the involvement of non-canonical, ligand-independent integrin signaling ^33, 38^. We generated a ITGB2 mutant (*mutITGB2*) that is ligand-binding-deficient, because aspartic acid 134 in the ligand-binding domain was substituted by alanine (Figure S6A). Overexpression of *mutITGB2* in A549 cells induced pEGFR and pMAPK, as well as increased the levels of VIM and ACTA2 in Galectin-3 (GAL3) -dependent manner (Figure 1F and S6B), demonstrating the involvement of non-canonical, ligand-independent integrin signaling during the phosphorylation-dependent activation of EGFR and MAPK. In addition, we observed co-localization of ITGB2 and pEGFR in NCI-H196 cells (Figure 1G, top), whereas ITGB6 and EGFR co-localized in A549 cells (Figure 1G, bottom). In summary, our results demonstrate that non-canonical ITGB2 signaling activates EGFR in SCLC.

### ITGB2 induces a novel SCLC gene expression signature

We retrieved and analyzed RNA-seq data of a previously published cohort of SCLC patients from the European Genome-Phenome Archive (Figure S7) ^39^. Correlation analysis of these RNA-seq based transcriptomes allowed us to group the SCLC patients into two clusters (C1 and C2; Figure 2A). Over-representation analysis (ORA) based on the Reactome database ^40^ for genes with increased expression in C2 (Figure 2B; 5,149 transcripts with fold change (FC) ≥ 3; Source Data S1) revealed a significant enrichment of genes related to the term “integrin cell surface interactions” (*P*=9.1E-3) as one of the top items of the ranked list. In addition, gene set enrichment analysis (GSEA) of the up-regulated transcripts in C2 (Figure 2C) showed a high enrichment score (ES) of 0.88 for “integrin cell surface interactions” (*P*=9E-3). Interestingly, lung tissue from SCLC patients in C2 showed significantly higher *ITGB2* expression (M=0.04; IQR=0.03) than the lung tissue from SCLC patients in C1 (M=0.01; IQR=0.01; *P*=2E-3; Figure 2D; Source Data S1). Moreover, we identified 93 transcripts that were upregulated in C2, in the SCLC cell line NCI-H196, as well as in A549 cells transiently transfected with *ITGB2* or *mutITGB2* (Figure 2E; Table S4). We named the set of 93 genes coding for these common upregulated transcripts as SCLC-ITGB2 gene expression signature (SCLC-ITGB2-sig). Comparing the enrichment of the genes grouped into the Kyoto Encyclopedia of Genes and Genomes (KEGG) term “SCLC” and the SCLC-ITGB2-sig (Figure S8), we observed ES of 0.52 (*P*=0.37) and 0.52 (*P*=0.36) for the KEGG-SCLC set in SCLC patients of C2 and C1, respectively, whereas the ES of the SCLC-ITGB2-sig significantly improved to 0.97 (*P*=8E-3) and 0.98 (*P*=8E-3) in SCLC patients of C2 and C1, respectively. Moreover, an external validation using independent RNA-seq data from SCLC cell lines ^41^ (Figure 2F) confirmed the improvement of the SCLC-ITGB2-sig (ES=0.77; FDR=0.50) as compared to the KEGG-SCLC set (ES=0.42; FDR=0.82). Thus, we propose the use of SCLC-ITGB2-sig for the identification of gene expression signatures related to SCLC. As shown in the heatmap in Figure 2G, the expression levels of the 93 genes, constituting the SCLC-ITGB2-sig, differentiate the two clusters of SCLC patients. Further, implementing data from 673 LC patients retrieved from Kaplan-Meier plotter ^42^ (Figure 2H) we found that lung cancer patients with increased levels of these 93 transcripts showed a significant lower overall survival of 88.7 months (*n*=372; HR=1.38; *P*=0.01) as compared to the overall survival of 127 months of patients with low expression levels (*n*=300). These findings support the clinical relevance of the SCLC-ITGB2-sig not only for stratification of SCLC patients, which might help to develop patient-tailored therapies, but also for prognosis prediction of LC patients.

**Figure 2:**
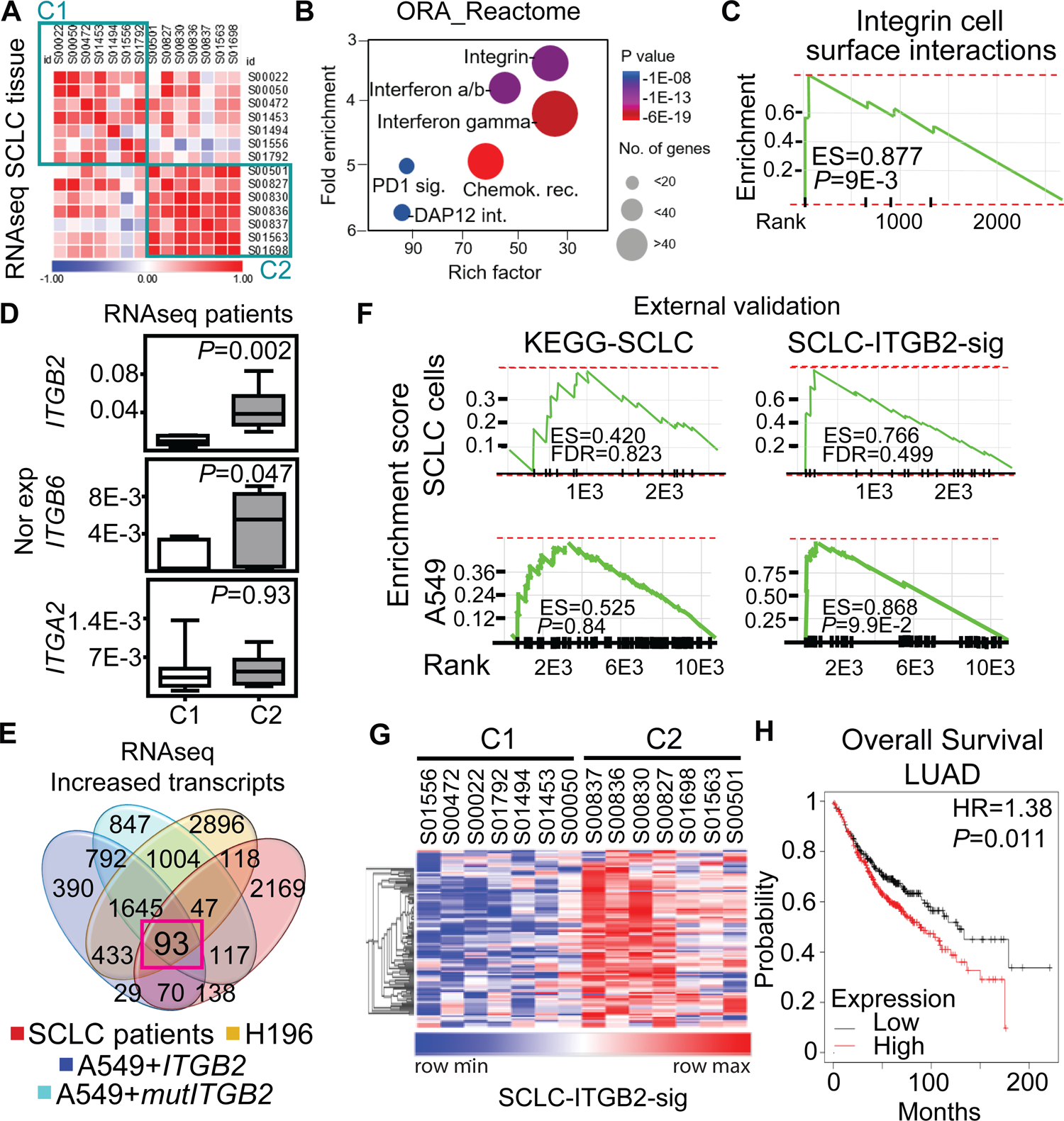
ITGB2 induces a novel SCLC gene expression signature. (**A**) Correlogram showing the correlation score matrix (RPKM values, Spearman correlation coefficient) across RNA-seq data of lung tissue of SCLC patients from the European Genome Archive (EGAS00001000299). SCLC patients were grouped in cluster 1 (C1) and cluster 2 (C2). (**B**) Bubble plot of top six enrichment of Reactome pathways in C2 by Overrepresentation Analysis (ORA). *P*-values after two-tailed t-test are shown by different color, the size of bubble indicate the gene count of each pathway. Sig., signaling; int., interactions; chemok., chemokine; rec., receptor. (**C**) Gene Set Enrichment Analysis (GSEA) using the fold change of genes inside the integrin pathway in B. ES, enrichment score; *P-*value after two-tailed t-test. (**D**) Box plots of RNA-seq-based expression analysis of indicated transcripts in SCLC patients C1 (*n*=7) and C2 (*n*=7). Values are represented as log2 RPKM. Nor exp, normalized expression to *GAPDH*. Box plots indicate median (middle line), 25th, 75th percentile (box) and 5th and 95th percentile (whiskers); *P*-values after two-tailed t-test. Source data are provided as Source Data S1. (**E**) Venn diagram comparing transcripts that were significantly increased in SCLC patients C2 compared to C1 (FC ≥ 3; *P* ≤ 0.05 after two-tailed t-test), NCI-H196 cells, A549 cells transfected either with *ITGB2* or *mutITGB2* (for all 3 cell lines, coding transcripts; FC ≥ 1.15; *P* ≤ 0.05 after two-tailed t-test) highlights a group of 93 transcripts that are common in all four groups, the SCLC-ITGB2 gene expression signature (SCLC-ITGB2-sig). See also Table S4. (**F**) External validation of the SCLC-ITGB2-sig. GSEA using independent RNA-seq data from SCLC cell lines ^41^ comparing the conventional SCLC signature in KEGG (left) versus the SCLC-ITGB2-sig (right) identified in E. ES, enrichment score; FDR, false discovery rate. (**G**) Hierarchical heatmap using RPKM of all 93 IDs of the SCLC-ITGB2-sig comparing SCLC patients in C1 to C2. Hierarchical clustering was performed using Person’s correlation based distance and average linkage. (**H**) Overall survival rates by Kaplan-Meier plotter of LUAD patients expressing low (*n*=300) or high (*n*=372) SCLC-ITGB2-sig (127 vs 88.7 months, respectively, *P*=0.011 after two-tailed t-test). HR, hazard ratio. See also Figure S7 and S8.

### Non-canonical ITGB2 signaling activates RAS/MAPK/ERK signaling

To further characterize the SCLC-ITGB2-sig, we performed a gene ontology (GO) enrichment analysis based on Biological Processes and found significant enrichment of genes related to GO terms involved in extracellular signal-regulated kinase 1/2 (ERK1/2) signaling pathway (Figure 3A). In addition, we also detected significant enrichment of genes related to RAS pathway (*P*=2.2E-2; FDR=0.26) and MAPK pathway (*P*=2.9E-3; FDR=0.1) (Figure 3B). These findings correlated with our previous results (Figure 1E-G and S5-S6), as Ras family proteins are known to be activated by signaling through EGFR ^43^. To further investigate the involvement of active RAS/MAPK/ERK signaling in SCLC, we performed RAS activation assay using whole cell protein extracts from A549, NCI-H196 and NCI-H82 cells (Figure 3C). This assay is based on the principle that the RAS binding domain (RBD) of the RAF kinase, one of the downstream RAS effector proteins, binds specifically to the GTP-bound form of RAS ^44^. Using RAF-RBD coated beads, we pulled down active, GTP-bound KRAS from protein extracts of NCI-H196 and NCI-H82 cells, supporting that RAS/MAPK/ERK signaling is active in SCLC. Interestingly, transfection of *ITGB2* or *mutITGB2* into A549 cells (Figure 3D) induced phosphorylation of EGFR, MAPK, RAF1 and ERK without significantly affecting the total levels of these proteins, thereby showing that non-canonical ITGB2 activates the EGFR and the RAS/MAPK/ERK signaling pathway. Further, we determined whether ITGB2 and KRAS are required for the intrinsic levels of pEGFR observed in both SCLC cell lines NCI-H82 and NCI-H196 (Figure 1E). We analyzed by WB protein extracts from NCI-H82 and NCI-H196 cells that were transfected with Ctrl, *ITGB2*- or *KRAS*-specific small interfering RNAs (siRNA, *siCtrl*, *siITGB2* or *siKRAS*; Figure 3E). Specific and efficient siRNA-mediated *ITGB2* or *KRAS* loss-of-function (LOF) reduced the levels of pEGFR and the downstream target of EGFR signaling VIM, demonstrating the requirement of ITGB2 and KRAS for the activation of EGFR and the RAS/MAPK/ERK signaling in SCLC. Interestingly, *siITGB2* or *siKRAS* transfection increased ITGB6 levels in both SCLC cell lines, supporting the mutual exclusive function of ITGB2 and ITGB6 shown in Figure 1 and S4.

**Figure 3:**
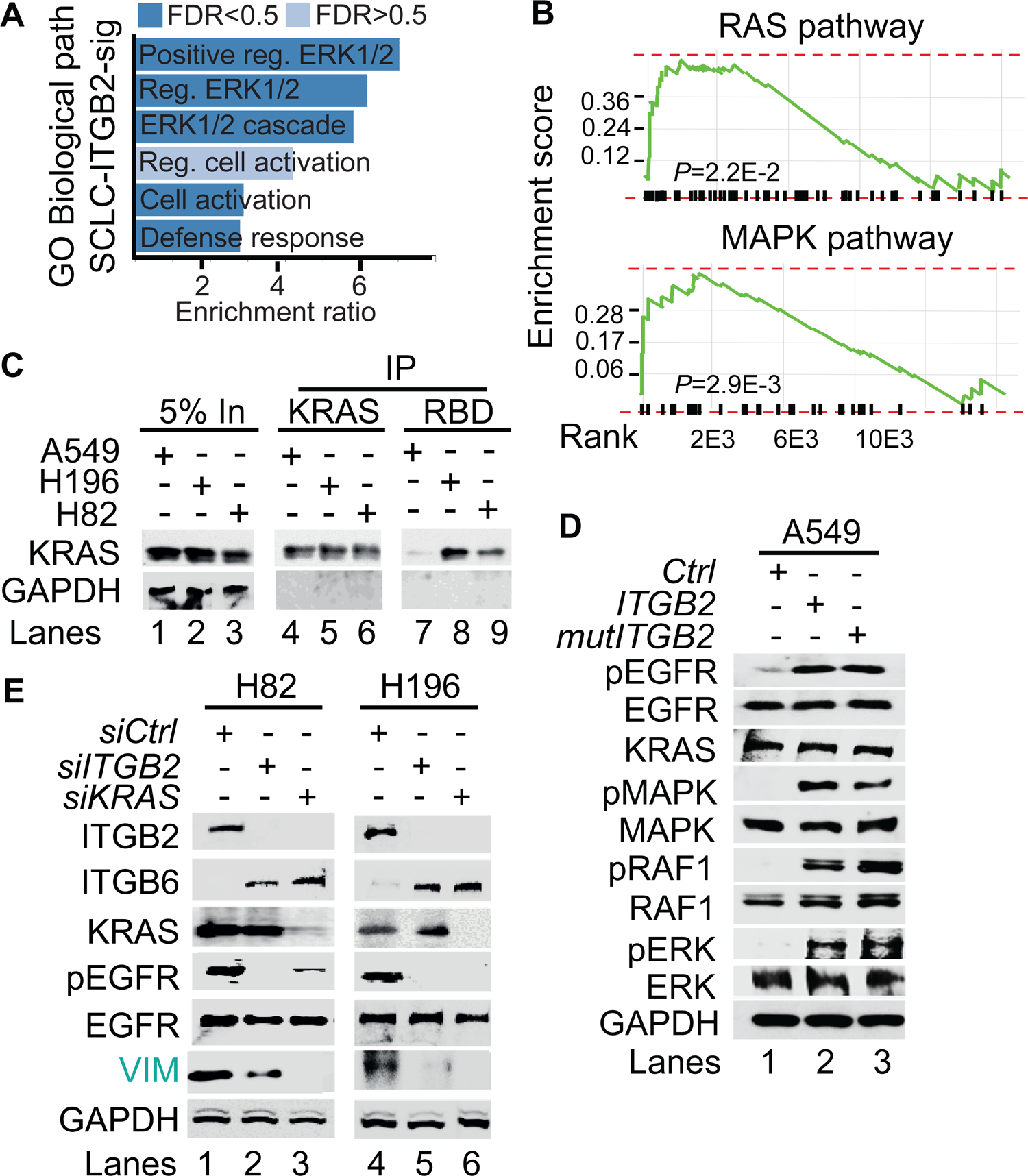
Non-canonical ITGB2 signaling activates RAS/MAPK/ERK signaling. (**A**) Gene Ontology (GO)-based enrichment analysis of biological pathways in the 93 IDs of the SCLC-ITGB2 gene expression signature (SCLC-ITGB2-sig) from Figure 2E using Webgestalt bioinformatics tool and plotted by highest enrichment ratio. Reg., regulation. (**B**) GSEA line profiles of SCLC-ITGB2-sig in RAS (Panther) and MAPK signaling pathways (KEGG). *P*-values after two-tailed t-test. (**C**) RAS activation assay. Protein extracts of A549, NCI-H196 and NCI-H82 were immunoprecipitated (IP) using a KRAS-specific antibody (KRAS) or RAF-RBD (RBD, active KRAS) coated beads. Co-IP proteins were analyzed by WB using the indicated antibodies. In, input, 5% of material used for the IP. (**D**) Total protein extracts of A549 cells transfected with *ITGB2* or ligand-binding-deficient D134A ITGB2 mutant (*mutITGB2*) were analyzed by WB using the indicated antibodies. pEGFR, active phosphorylated epidermal growth factor receptor; pMAPK, phosphorylated mitogen-activated protein kinase; pRAF1, phosphorylated proto-oncogene serine/threonine-protein kinase; pERK, phosphorylated extracellular signal-regulated kinase. (**E**) Total protein extracts of NCI-H82 and NCI-H196 cells transfected with small interfering RNA specific for *ITGB2* (*siITGB2*) or *KRAS* (*siKRAS*) were analyzed by WB using the indicated antibodies. Vimentin (VIM) as product of a downstream gene target of EGF signaling is highlighted in green.

To gain further insight into the mechanism of ITGB2-induced activation of EGFR, we performed Co-IP using ITGB6- or ITGB2-specific antibodies in protein extracts from A549 cells transiently transfected with *Ctrl* (empty vector), *ITGB6* or *ITGB2* (Figure 4A). We found that endogenous ITGB6 interacts with non-phosphorylated EGFR and *ITGB2* overexpression abolished this interaction. Further, *ITGB2* transfection induced pEGFR and reduced ITGB6 levels, confirming our previous results (Figure 1E and S4-S6). Moreover, over-expressed ITGB2 co-precipitated pEGFR demonstrating the interaction of ITGB2 with the active form of this receptor. These results were complemented by Co-IP experiments using protein extracts from NCI-H196 cells transiently transfected with *Ctrl* or *ITGB6* (Figure 4B). We detected interaction of endogenous ITGB2 with pEGFR, confirming the results in A549 cells after overexpression of *ITGB2*. Further, *ITGB6* overexpression abolished the ITGB2-pEGFR-interaction. Our results demonstrate the interaction between endogenous ITGB6 and EGFR in the NSCLC cell line A549, whereas endogenous ITGB2 interacts with pEGFR in the SCLC cell line NCI-H196. Moreover, these interactions appear mutually exclusive, since overexpression of *ITGB2* or *ITGB6* abolished the interaction of the complementary integrin receptor beta subunit with EGFR. Remarkably, precipitation of endogenous EGFR in NCI-H196 cells confirmed the pEGFR-ITGB2 interaction (Figure 4C). Moreover, siRNA-mediated LOF of *ITGB2*, *GAL3* or *KRAS* abolished the pEGFR-ITGB2 interaction, thereby demonstrating the specificity of the pEGFR-ITGB2 interaction and the requirement of GAL3 and KRAS for this interaction. Further Co-IP experiments in A549 cells (Figure 4D) showed that endogenous KRAS interacted with overexpressed ITGB2, mutITGB2, endogenous EGFR and pEGFR and GAL3 in a GAL3-dependent manner (lanes 5 to 8). Interestingly, pull down of active, GTP-bound KRAS using RAF-RBD coated beads (lanes 9 to 12) co-precipitated overexpressed ITGB2, mutITGB2, GAL3 and endogenous EGFR, but not pEGFR, in a GAL3-dependent manner. Our results support that the KRAS fraction interacting with pEGFR is inactive (compare lanes 5 to 8 with 9 to 12). The model in Figure 4E summarizes our results. Endogenous ITGB6 interacts with EGFR in the NSCLC cell line A549 (Figure 4E, left). The ITGB6-EGFR-interaction is abolished upon *ITGB2* or *mutITGB2* transfection, suggesting a functional switching of a complex containing ITGB6-EGFR in NSCLC to a complex containing ITGB2-EGFR in SCLC. Supporting this line of ideas, over-expressed *ITGB2* or *mutITGB2* in A549 cells, or endogenous ITGB2 in the SCLC cell line NCI-H196 interact with endogenous pEGFR (Figure 4E, middle). However, our results indicate the formation of a multimeric protein complex in at least two different forms. One form contains ITGB2, pEGFR, GAL3 and inactive, GDP-bound KRAS (Figure 4E, middle). The other form contains ITGB2, EGFR, GAL3 and active, GTP-bound KRAS (Figure 4E, right). Considering the results presented in Figure 1 and 3, we propose that these forms of the multimeric protein complex occur in sequential order during non-canonical ITGB2-mediated activation of KRAS/MAPK/ERK signaling in SCLC.

**Figure 4:**
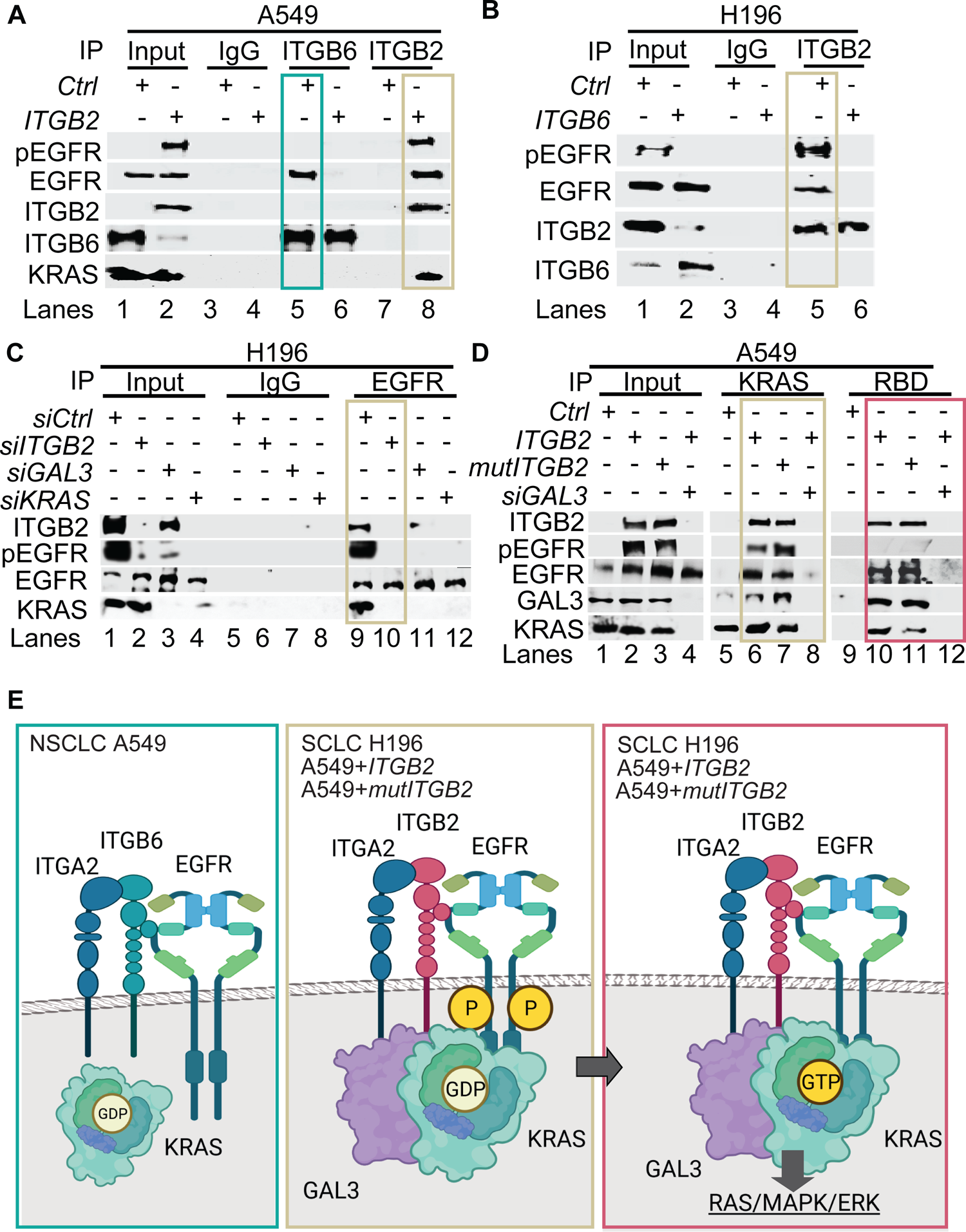
Different multimeric protein complexes sequentially occur during non-canonical ITGB2-mediated activation of KRAS/MAPK/ERK signaling in SCLC. (**A**) Total protein extracts of A549 cells transfected with empty vector (*Ctrl*) or *ITGB2* were immunoprecipitated (IP) using either immunoglobulin G (IgG, as control) or ITGB6 and ITGB2-specific antibodies. Co-IP proteins were analyzed by WB using the indicated antibodies. Input, 5% of material used for the IP. Squares indicate conditions in which endogenous ITGB6 interacts with inactive EGFR (green) and overexpressed ITGB2 interacts with active pEGFR (gold). (**B**) Total protein extracts of NCI-H196 cells transfected with empty vector (*Ctrl*) or *ITGB6* were immunoprecipitated (IP) using either immunoglobulin G (IgG, as control) or ITGB2-specific antibodies. Co-IP proteins were analyzed by WB using the indicated antibodies. Input, 5% of material used for the IP. Gold square indicates conditions in which endogenous ITGB2 interacts with endogenous, active pEGFR. (**C**) Protein extracts of NCI-H196 cells transfected with small interfering RNA specific for *ITGB2* (*siITGB2*), *GAL3* (*siGAL3*) or *KRAS* (*siKRAS*) were immunoprecipitated (IP) using either immunoglobulin G (IgG, as control) or EGFR-specific antibodies. Co-IP proteins were analyzed by WB using the indicated antibodies. In, input, 5% of material used for the IP. Gold square indicates conditions showing the ITGB2-pEGFR-interaction is specific and GAL3- and KRAS-dependent. (**D**) RAS activation assay. Protein extracts of A549 cells transfected with *ITGB2* or ligand-binding-deficient D134A ITGB2 mutant (*mutITGB2*) and *siGAL3* were immunoprecipitated (IP) using KRAS-specific antibody (KRAS) or RAF-RBD (RBD, active KRAS) coated beads. Co-IP proteins were analyzed by WB using the indicated antibodies. In, input, 5% of material used for the IP. Gold square highlights conditions in which KRAS interacts with ITGB2, mutITGB2, GAL3, EGFR and pEGFR in GAL3-dependent manner. Magenta square highlights conditions in which active, GTP-bound KRAS interacts with ITGB2, mutITGB2, GAL3 and EGFR, but not with pEGFR. (**E**) Model. Left, endogenous ITGB6 interacts with EGFR in the NSCLC cell line A549. Middle, endogenous ITGB2 interacts with endogenous pEGFR in the SCLC cell line NCI-H196, or in A549 cells after *ITGB2* or *mutITGB2* transfection. Further, results from RAS activation assays indicate the formation of a multimeric protein complex in two different forms, one form containing ITGB2, pEGFR, GAL3 and inactive, GDP-bound KRAS (middle) and the other form containing ITGB2, EGFR, GAL3 and active, GTP-bound KRAS (right), both forms occurring in sequential order during non-canonical ITGB2-mediated activation of KRAS/MAPK/ERK signaling in SCLC.

### Extracellular vesicles containing ITGB2 activate RAS/MAPK/ERK signaling and induce SCLC proteins

ORA of the SCLC-ITGB2-sig based on the Reactome database revealed significant enrichment of genes related to glycosphingolipid metabolism (Figure 5A). In addition, we found elevated expression of the proto-oncogenes *MYCN* and *MYCL* in SCLC cell lines (Figure 5B). Furthermore, GO enrichment analysis based on Biological Processes of transcripts with high levels in SCLC cell lines revealed significant enrichment of genes related to the GO term “Vesicle transport” (Figure S9A-B). Since all these events are related to the secretion of EVs ^45^, we isolated EVs from the cell culture medium, in which A549, NCI-H82 and NCI-H196 cells were grown, and characterized them by various downstream analyses (Figure 5C and S9C). EVs produced by these three cell lines showed similar size with a radius of approximately 45 nm (Figure S9C). Further, high-resolution mass spectrometry analysis of the protein cargo of the isolated EVs revealed 189 proteins that were common in EVs from NCI-H196 cells transfected with control plasmid (*Ctrl*) and from A549 cells transfected either with *ITGB2* or *mutITGB2* (Figure 5D; Table S5 and Figure S9D). Panther-based ORA of these common 189 proteins (Figure 5E) showed significant enrichment of proteins involved in “Ras pathway” and “Integrins signaling pathway”, correlating with our previous results (Figure 1-4). Further, WB of the protein cargo of EVs produced by A549 cells that were transiently transfected with *Ctrl*, *ITGB2* or *mutITGB2* (Figure 5F) showed similar levels of ITGA2, as well as of the EVs markers TSG101 (tumor susceptibility gene 101 protein) and the CD63 antigen ^46, 47^, whereas MYCN was specifically detected after *ITGB2* or *mutITGB2* transfection, confirming our mass spectrometry results (Table S5). Interestingly, confocal microscopy in sections of hPCLS that were incubated with EVs produced by A549 cells revealed ITGB2 staining when the A549 cells were transfected with *ITGB2* prior EVs isolation (Figure 5G), supporting that EVs influence the gene expression signature of the treated hPCLS. Importantly, this interpretation was confirmed by WB of protein extracts from hPCLS incubated with A549 EVs (Figure 5H). Protein extracts of hPCLS showed increased ITGB2 levels when they were incubated with EVs produced by A549 cells that were transfected with *ITGB2* or *mutITGB2*. Interestingly, EVs produced by A549 cells transfected with *ITGB2* or *mutITGB2* induced phosphorylation-dependent activation of EGFR and MAPK, as well as increased levels of the downstream targets of EGFR signaling VIM and FN1, demonstrating activation of the RAS/MAPK/ERK signaling pathway in the treated hPCLS. Moreover, EVs isolated from *ITGB2*- or *mutITGB2*-transfected A549 cells induced NKX2-1, EZH2, ASH-1 and MYCN in treated hPCLS, whereas TP53 levels were reduced, thereby mimicking gene expression patterns that are characteristic of SCLC ^48, 49^. As we have previously shown that the ribonuclease (RNase) binase inhibits oncogenic KRAS ^44^, we included binase treatment in our experimental setting. Remarkably, binase treatment counteracted the effects caused by EVs isolated from *ITGB2*- or *mutITGB2*-transfected A549 cells, thereby demonstrating the causal involvement of KRAS activation.

**Figure 5:**
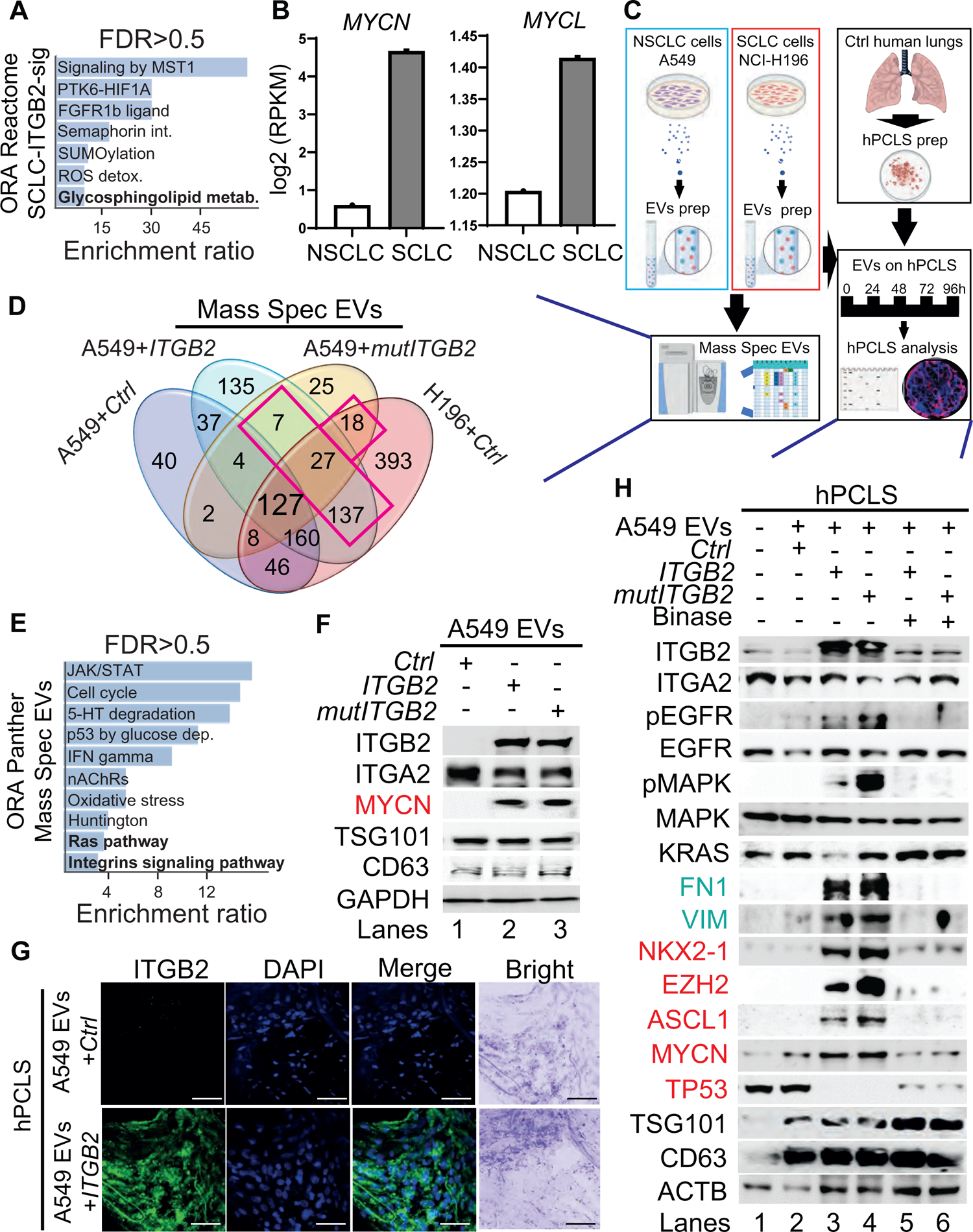
Extracellular vesicles containing ITGB2 activate RAS/MAPK/ERK signaling and induce SCLC proteins. (**A**) Reactome-based enrichment analysis of significant pathways in the 93 IDs of the SCLC-ITGB2 gene expression signature (SCLC-ITGB2-sig) from Figure 2E using Webgestalt bioinformatics tool and plotted by highest enrichment ratio. Int., interactions; metab., metabolism. (**B**) RNA-seq-based expression analysis of indicated transcripts in non-small cell lung cancer (NSCLC; *n*=33) and small cell lung cancer (SCLC; *n*=17) cell lines. Values were normalized to *GAPDH* and represented as log2 of reads per kilobase per million (RPKM). Bar plots show data as means; error bars, s.e.m. (**C**) Scheme of experiments with EVs isolated from the cell culture medium of NSCLC and SCLC cell lines. Characterization of the protein cargo of the isolated EVs by high-resolution mass spectrometry (HRMS) analysis. Characterization of human precision-cut lung slices (hPCLS) that were treated with isolated EVs. (**D**) Venn diagram comparing proteins that were detected by HRMS in EVs from control transfected A549 cells, NCI-H196 cells, as well as from A549 cells transfected either with *ITGB2* or *mutITGB2* highlights a group of 189 proteins that are common for the last 3 conditions. See also Table S5. (**E**) Panther-based enrichment analysis of significant pathways in the 189 proteins highlighted in D, using Webgestalt bioinformatics tool and plotted by highest significance enrichment ratio. (**F**) Total protein extracts of EVs from A549 cells transfected with *ITGB2* or *mutITGB2* were analyzed by WB using the indicated antibodies. (**G**) Confocal microscopy after immunostaining with specific antibodies against ITGB2 in hPCLS incubated with EVs from A549 cells previously transfected with *Ctrl* or *ITGB2*. DAPI, nucleus. Scale bars, 500 μm. (**H**) Total protein extracts of hPCLS incubated with EVs from A549 cells previously transfected with *ITGB2* or *mutITGB2* alone or in combination with binase were analyzed by WB using the indicated antibodies. Products of downstream gene targets of EGF signaling (green) and SCLC proteins (red) are highlighted. See also Figure S9 and S10.

### ITGB2 loss-of-function and binase inhibit SCLC-associated proteins

Since EVs produced by *ITGB2*- or *mutITGB2*-transfected A549 cells were able to induce RAS/MAPK/ERK signaling and SLCL markers in hPCLS, we decided to investigate the EVs produced by the SCLC cell line NCI-H196 either non-transfected or transfected with *siCtrl* or *siITGB2* or treated with binase (Figure 6). WB of the protein cargo of EVs produced by non-transfected or *siCtrl*-transfected NCI-H196 cells showed similar levels of ITGB2, MYCN, TSG101 and CD63 (Figure 6A), whereas *siITGB2* transfection or binase treatment of the NCI-H196 cells prior EVs isolation reduced the ITGB2 and MYCN levels in the isolated EVs. Further, we analyzed the effects caused on hPCLS by EVs produced by NCI-H196 cells under the four conditions specified above (Figure 6B-D). EVs produced by *siCtrl*-transfected NCI-H196 cells increased cell proliferation and cell number of hPCLS after 96h treatment (Figure 6B). We also detected ITGB2 and VIM by confocal microscopy in sections of hPCLS that were incubated with EVs produced by NCI-H196 cells (Figure 6C), supporting that such EVs induced in the treated hPCLS gene expression patterns that are similar to SCLC. These results were complemented by WB of protein extracts from hPCLS incubated with NCI-H196 EVs (Figure 6D). EVs produced by NCI-H196 cells induced in hPCLS ITGB2, phosphorylation-dependent activation of EGFR and MAPK, as well as increased levels of the downstream targets of EGFR signaling VIM and FN1, demonstrating activation of the RAS/MAPK/ERK signaling pathway in the treated hPCLS. Moreover, EVs produced by NCI-H196 cells induced EZH2, H3K27me3 (Histone 3 tri-methylated at lysine 27) and NKX2-1 in treated hPCLS, thereby mimicking gene expression patterns that are characteristic of SCLC ^48, 49^. Remarkably, the effects induced by NCI-H196 EVs on hPCLS were counteracted by *siITGB2* transfection or binase treatment of NCI-H196 cells prior EVs isolation (Figure 6B-D). These findings are in line with the reducing effect of binase on cancer hallmarks, such as colony formation and cell viability (Figure S10A-C), suggesting both, ITGB2-LOF as well as binase treatment, for the development of therapies against SCLC.

**Figure 6:**
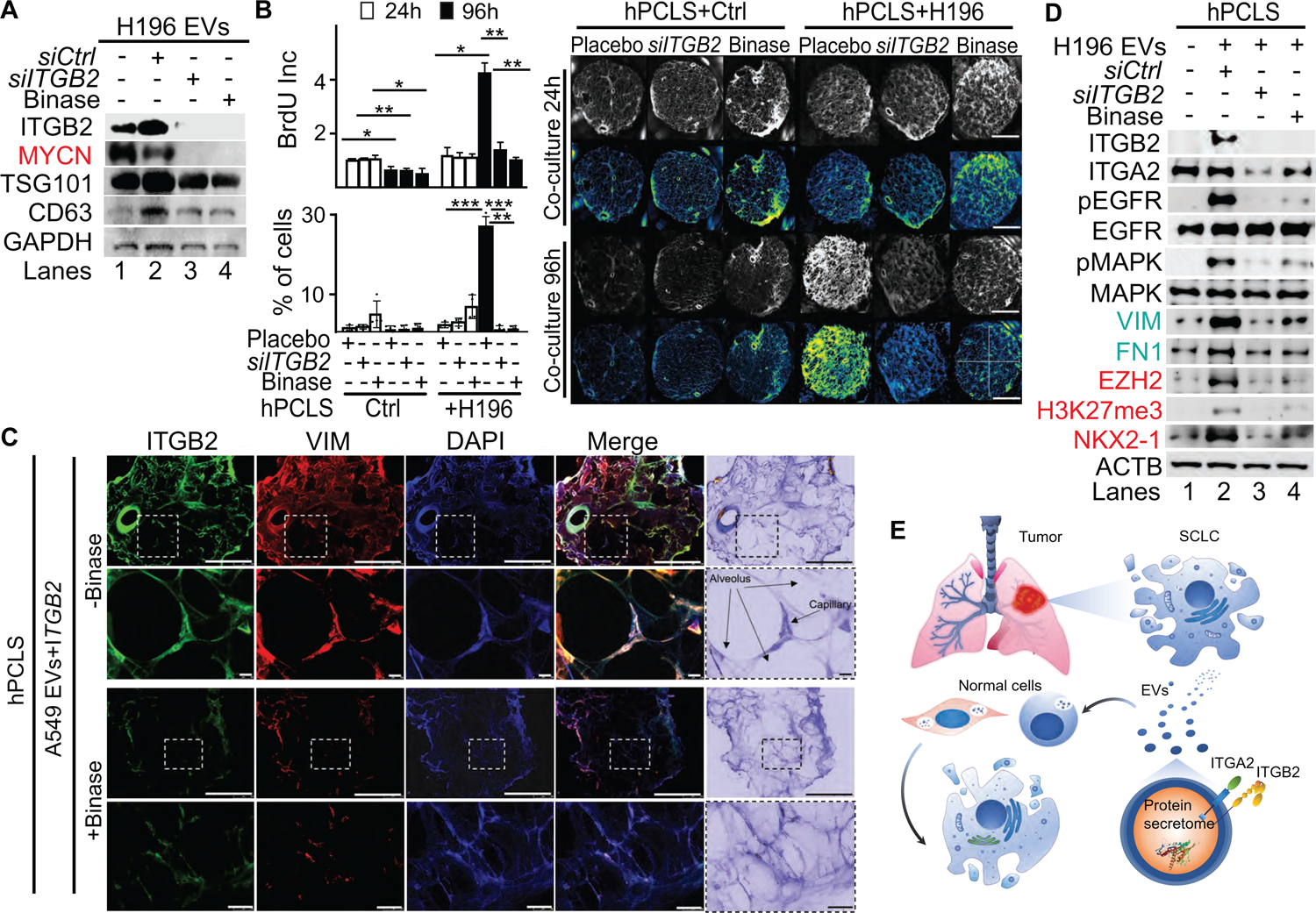
ITGB2 loss-of-function and binase inhibit SCLC. (**A**) Total protein extracts of EVs from NCI-H196 cells transfected with control or *ITGB2*-specific small interfering RNA (*siCtrl* or *siITGB2*), or treated with binase, were analyzed by WB using the indicated antibodies. (**B**) Left, cell proliferation of hPCLS co-cultured without or with NCI-H196 cells previously transfected with *siITGB2* or treated with binase was measured by the colorimetric method using BrdU incorporation (top) cell number quantification (bottom). Data are shown as means ± s.e.m (*n*=3 independent experiments); asterisks, *P*-values after one tailed t-test, \*\*\**P*˂0.001; \*\**P*˂0.01; \**P*˂0.05. Right, representative live microscopy images to the bar plots. Quadrants used for quantification are indicated. Scale bars, 500 μm (**C**) Confocal microscopy after immunostaining with specific antibodies against ITGB2 and VIM in hPCLS incubated with EVs from A549 cells previously transfected with *ITGB2* and non-treated or treated with binase. DAPI, nucleus. Scale bars, 500 μm. Squares are respectively shown below at higher magnification. (**D**) Total protein extracts of hPCLS incubated with EVs from NCI-H196 cells previously transfected with *siCtrl* or *siITGB2* alone or in combination with binase were analyzed by WB using the indicated antibodies. Products of downstream gene targets of EGF signaling (green) and SCLC proteins (red) are highlighted. (**E**) Model. In SCLC, high ITGB2 induces a KRAS-driven secretory phenotype of ITGB2/ITGA2 loaded EVs, which total protein cargo induces a SCLC-like phenotypic transformation in normal cells. See also Figure S9 and S10.

## Discussion

In the present study, we showed that ITGB6 interacts with the inactive EGFR in NSCLC, whereas in SCLC, ITGB2 reduces the levels of ITGB6, and interacts with and activates EGFR. In addition, we demonstrated that the EVs produced by a SCLC cell line induced in hPCLS both, non-canonical ITGB2-mediated activation of KRAS/MAPK/ERK signaling and SCLC proteins, supporting the hypothesis that the cargo of EVs may influence the gene expression signature of hPCLS (Figure 6D). Following this line of ideas, siRNA-mediated *ITGB2*-LOF or binase treatment of the SCLC cell line led to the production of EVs that lost the capacity to induce KRAS/MAPK/ERK signaling and SCLC proteins in hPCLS. On the one hand, our results provide a model (Figure 6E) to study the effects of EVs on specific tissues by targeted cargo modification through manipulation of the gene expression in EV-producing cells. On the other hand, they also provide a plausible explanation for the resistance of SCLC to tyrosine kinase inhibitors targeting EGFR (EGFR-TKIs), including Erlotinib, Gefitinib and Afatinib, and suggest the development of novel therapeutic strategies against SCLC combining EGFR-TKIs and ITGB2-LOF. Alternatively, treating the lung of SCLC patients with EVs produced by NSCLC cell lines to achieve lung tissue sensitivity to EGFR-TKIs may be one treatment strategy worthy of exploration. Furthermore, determining high levels of ITGB2 in cases of NSCLC showing enhanced EGFR signaling in the absence of somatic mutations may validate ITGB2 as a promising therapeutic target in NSCLC. This reasoning is supported by the current use of EGFR-TKIs in lung adenocarcinoma patients with EGFR mutations ^50, 51^. The majority of these hyperactive EGFR mutants harbor either a point mutation, in which leucine 858 is substituted by arginine (L858R), or a deletion involving 5 codons coding for the amino acids at the positions 746-750 (ΔEx19) ^52^. However, a secondary point mutation in the EGFR kinase domain that substitutes threonine 790 by methionine (T790M) produces drug-resistant EGFR variants, which are present in 50% of patients that developed resistance to EGFR-TKIs treatment ^53^. Surprisingly, studies of biopsies have shown that 5-15% NSCLC patients undergo histological transformation to SCLC upon acquisition of therapy resistance ^54^. It will be extremely interesting to determine whether this histological transformation from NSCLC to SCLC upon acquired EGFR-inhibitor resistance is produced by a functional switch from ITGB6-ITGA2 to ITGB2-ITGA2 during EGFR complex formation.

Various aspects of the signaling model proposed here might be applicable in a broader context. Specific integrin receptor subunits have been identified as biological markers and potential therapy targets to tumor progression and metastasis in a wide range of cancers including glioblastoma, pancreatic carcinomas, breast cancer, leukemia, among others ^33, 55^. In addition, recent reports demonstrated functional correlation between the switch of specific integrin subunits and an aggressive phenotype of cancer cells. For instance, ITGA2-ITGB1 promotes chemotherapy resistance of T-cell acute lymphoblastic leukemia ^56^. Further, ITGB1 has been reported to trigger an EGFR ligand-independent proliferation signaling in pancreatic ductal adenocarcinoma, bypassing the EGFR-blocking effect of the anti-EGFR monoclonal antibody, Cetuximab ^57^. These reports suggest that targeting specific integrin subunits might be beneficial against a wider spectrum of cancer types. Supporting this hypothesis, depletion of ITGB3, ITGB4 and ITGB5 reduced angiogenesis and tumor growth in breast cancer ^31^.

Another interesting aspect of our model is the functional competition between ITGA2-ITGB2 in SCLC and ITGA2-ITGB6 in NSCLC. Similar competitions have been observed between ITGAM-ITGB2 and ITGA5-ITGB1 or ITGAv-ITGB3 and ITGA5-ITGB1 heterodimers regulating migration or trafficking of leukocytes ^58, 59^. Recently, a mutual competition between ITGA5-ITGB1 blocking EGF signaling and ITGA6-ITGB4 enhancing EGF signaling has been reported ^60^, highlighting the specific interaction between integrin subunits mediating different functions in EGF signaling. Interestingly, EGFR interacts in the cell membrane with glycosylated regulatory partners including proteoglycans, as syndecans ^61, 62^. Remarkably, *N*-glycosylation of specific domains in the ITGA5 subunit appears critical for different processes mediating its biological function, such as ITGA5-ITGB1 heterodimer formation, its expression on the cell surface, ligand binding, EGFR-ITGA5-ITGB1 complex formation and its inhibitory effect on EGFR ^60, 63, 64^. Similar to ITGA5, it has been also reported that *N*-glycosylation of ITGB4 is essential for appropriate EGFR-ITGB4-ITGA6 complex formation on the cell surface^65^. Future work will determine whether a similar mechanism of *N*-glycosylation participates in switching the EGFR complex formation from ITGB6-ITGA2 to ITGB2-ITGA2 during the histological transformation from NSCLC to SCLC observed upon acquired EGFR-TKIs resistance ^54^.

We uncovered a mechanism of non-canonical ITGB2-mediated EGFR activation that explains EGFR-TKIs resistance in SCLC cases lacking EGFR mutations. Our results not only support the use of ITGB2 and the newly identified SCLC-ITGB2-sig as diagnostic markers for SCLC, but also as targets to develop therapeutic strategies against this extremely aggressive type of LC.

## Materials and Methods

### Study population

The study was performed according to the principles set out in the WMA Declaration of Helsinki and the protocols approved by the institutional review board and ethical committee of Regional Hospital of High Specialties of Oaxaca (HRAEO), which belongs to the Ministry of Health in Mexico (HRAEO-CIC-CEI 006/13), the Medicine Faculty of the Justus Liebig University in Giessen, Germany (Ethical Votum 68/13) and the Hannover Medical School (no. 2701-2015). In this line, all patient and control materials were obtained through the HRAEO in Mexico, the Institute for Pathology of Justus Liebig University in Giessen and the Biobank from the Institute for Pathology of the Hannover Medical School as part of the BREATH Research Network. We used anonymized patient material.

Formalin-fixed paraffin-embedded (FFPE) human lung tissue samples of either diagnostic transbronchial or bronchial biopsies or oncologic resections were retrospectively collected. All cases were reviewed and staged by an expert panel of pulmonologist and oncologist. FFPE tissue samples of LC patients comprised approximately 80% tumor cells. The control population for the analysis included lung tissue that was taken from macroscopically healthy adjacent regions of the lung of LC patients. Corresponding clinical data for matched patients with LUAD (*n*=11) were obtained from The Genome Cancer Atlas (TCGA, tcga-data.nci.nih.gov/doc/publications/tcga/). Data are publicly available and open-access. Clinical characteristics of LUAD patients are presented in Table S3.

### Cell culture, transfection, treatment and siRNA-mediated knockdown

Mouse lung epithelial cells MLE-12 (ATCC CRL-2110) were cultured in complete DMEM/F12 (5% FCS, 1% Penn-strep) at 37^°^C in 5% CO_2_. Human SCLC cells NCI-H82 (ATCC HTB-175) and NCI-H196 (ATCC CRL-5823), and NSCLC cells A549 (ATCC CCL-185) were cultured in complete RPMI (10% FCS, 1% Penn-strep) at 37^0^C in 5% CO_2_. During subculturing, cells were 1x PBS washed, trypsinized with 0.25% (w/v) Trypsin and subcultivated at the ratio of 1:5 to 1:10. The cell lines used in this paper were mycoplasma free. They were regularly tested for mycoplasma contamination. In addition, they are not listed in the database of commonly misidentified cell lines maintained by ICLAC. Cells were transfected with plasmid DNA or siRNA using Lipofectamine 2000 (Invitrogen) following the manufacturer’s instructions, and harvested 48 h later for further analysis. *ITGA2-HIS* (Addgene, #51910), *ITGB2-YFP* (Addgene, #8638) and *ITGB6-GFP* (Addgene, #13293) mammalian expression constructs were used for respective gene overexpression in cell lines. *siITGB2* (EHU133911) was purchased from Sigma. Empty vectors and *siCtrl* were used as a negative control.

### Bacterial culture and cloning

For cloning experiments, chemically competent *E. coli* TOP10 (ThermoFisher Scientific) were used for plasmid transformation. TOP10 strains were grown in Luria broth (LB) at 37^0^C with shaking at 180rpm for 16h or on LB agar at 37^0^C overnight.

### RNA isolation, reverse transcription, quantitative PCR and TaqMan assay

Expression analysis by qRT-PCR were performed as previously described ^66^. Briefly, total RNA from cell lines was isolated using the RNeasy Mini kit (Qiagen) and quantified using a Nanodrop Spectrophotometer (ThermoFisher Scientific, Germany). Human lung tissue samples were obtained as FFPE tissues, and eight sections of 10 μm thickness were used for total RNA isolation using the RecoverAll Total Nucleic Acid Isolation Kit for FFPE (Ambion). Clinical characteristics of SCLC patients are presented in Table S2. Synthesis of cDNA was performed using 0.2-1 µg total RNA and the High Capacity cDNA Reverse Transcription kit (Applied Biosystems). Quantitative real-time PCR reactions were performed using SYBR^®^ Green on the Step One plus Real-time PCR system (Applied Biosystems). Housekeeping genes *HPRT* and *GAPDH* were used to normalize gene expression. Primer pairs used for gene expression analysis are described in the following table:

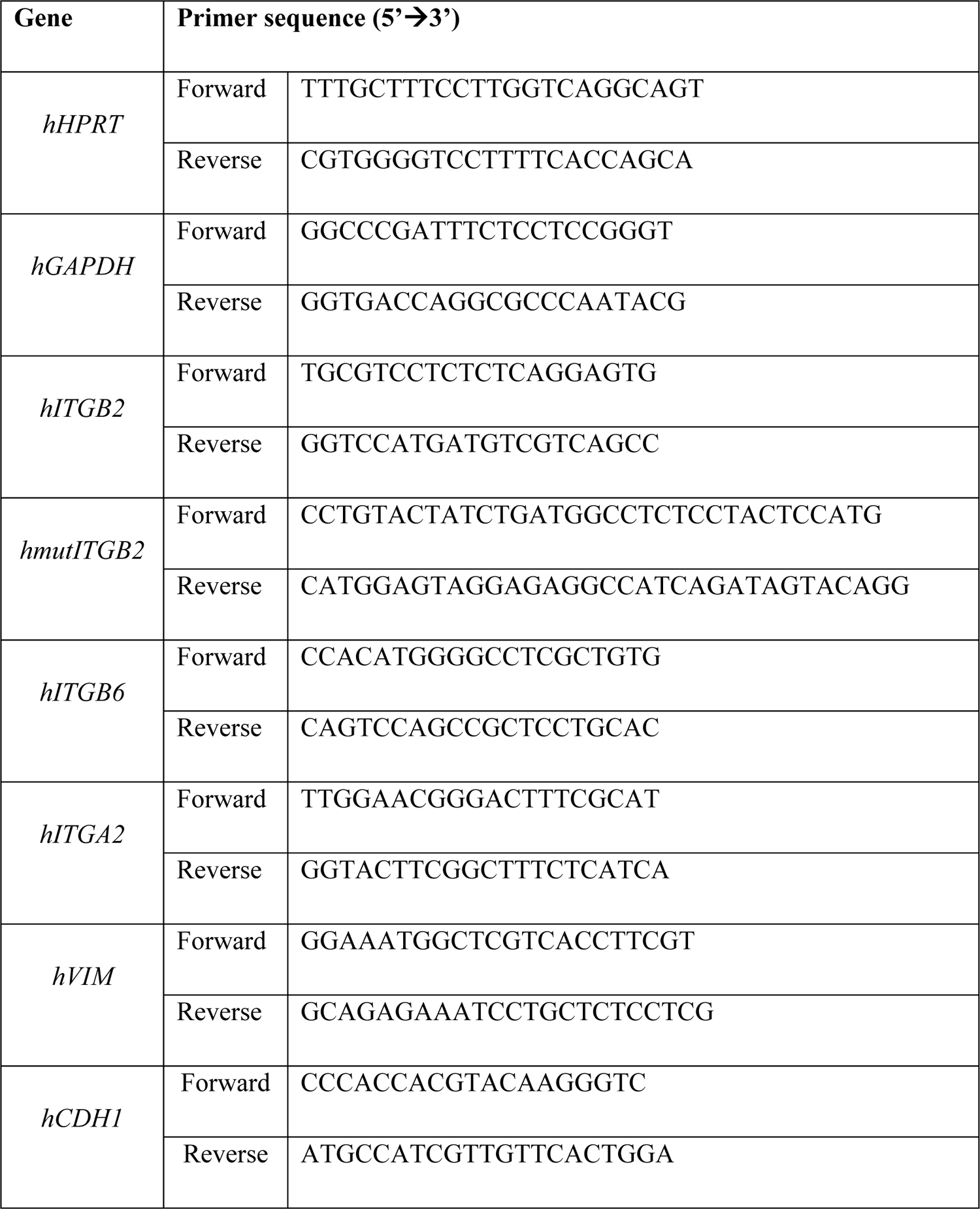

### Immunofluorescence and confocal microscopy

Immunostaining was performed as previously described ^67^. Briefly, cells were grown on coverslips, fixed with 4% PFA for 10min at RT and permeabilized with 0.4% Triton-X100/1xPBS for 10min at RT. For non-adherent cells, slides were previously coated with poly-L-lysine. During immunostaining procedure, all incubations and washes were performed with histobuffer containing 3% bovine serum albumin and 0.2% Triton X-100 in 1xPBS, pH7.4. Non-specific binding was blocked by incubating with donkey serum and histobuffer (1:1 (v/v) ratio) for 1h. Cells were then incubated with primary and then secondary antibodies for 1hr followed by nuclear staining. Immunostaining of cells were examined with a confocal microscope (Zeiss). Antibodies used were specific anti-ITGB2 (R&D Systems), anti-ITGB6 (R&D Systems), anti-GFP (Santa Cruz), anti-pEGFR (Cell Signaling), and anti-EGFR (Cell Signaling), were used. Alexa 488, Alexa 555 or Alexa 594 tagged secondary antibodies (Invitrogen, Germany, dilution 1:2000) were used. DAPI (Sigma, Germany) used as nuclear dye.

Paraffin-embedded lung tissue sections (3-μm thick) were deparaffinized in xylene and rehydrated in a graded ethanol series to PBS (pH 7.2). Antigen retrieval was performed by pressure cooking in citrate buffer (pH 6.0) for 15 min. Double immunofluorescence staining was performed with primary antibodies anti-ITGB2 (R&D Systems), anti-ITGB6 (R&D Systems), anti-GFP (Santa Cruz), anti-VIM (Cell Signaling) and anti-ASH1 (Chemicon) were used. After overnight incubation with specific primary antibodies, slides were washed and incubated with the respective secondary antibodies, Alexa 488–, Alexa 555– and Alexa 594-conjugated goat anti-rabbit IgG (dilution 1:1000, Molecular Probes, Eugene, OR) for 1h. All sections were counterstained with nuclear DAPI (1:1000) and mounted with fluorescent mounting medium (Dako).

### Co-immunoprecipitation (Co-IP) and Western blot

Total protein extracts from different cell lines were prepared in 1 ml ice cold RIPA buffer [(50 mM (pH7.5) Tris-HCl, 150mM NaCl, 1% Triton X-100 (Sigma), 0.5% Na-deoxycholate (Sigma), 0.1% SDS, 0.2 M imidazole (Sigma), 10 mM NaF (Sigma), 2 mM Na3VO4 (Sigma), 1 mM phenylmethylsulfonyl fluoride (PMSF) and protease inhibitor cocktail (Calbiochem)]. Detergent-insoluble material was precipitated by centrifugation at 14,000 rpm for 30 min at 4 °C. The supernatant was transferred to a fresh tube and stored at −20 °C. Protein concentration was estimated using Bradford assay, using serum albumin as standard. 5 μl of serial dilutions of standard protein and samples were mixed with 250 μl of Bradford reagent (500-0205, BIO-RAD Quick Start™). Samples were incubated 10 minutes at room temperature and measured at OD595 using an ELISA plate reader (TECAN Infinite M200 Pro). Co-IP was performed as described ^68^ with minor adaptations. For immunoprecipitation of membrane proteins, a total of 5 x 10^7^ cells were collected and washed three times in cold PBS, spun down at 270g for 10min at 4°C. Protein extracts were obtained as described above using cell immunoprecipitation (IP) buffer [(50 mM, pH 7.5) Tris-HCl, 1 mM MgCl_2_, 1 mM CaCl_2,_ 150 mM NaCl, 1% NP40, 1 mM phenylmethylsulfonyl fluoride (PMSF) and protease inhibitor (Calbiochem)] Protein concentrations were determined as described above. Precleared protein lysates (500 μg per immunoprecipitation) were resuspended in 500 μl IP buffer and incubated with the appropriate antibodies on ice for 2 h and then 30 μl protein-G-sepharose beads (GE Healthcare; equilibrated once in 10 ml water and three times in washing buffer) were added and incubated overnight at 4°C. Beads were collected and washed 5 times with 500 μl ice-cold washing buffer. 30 μl 2x SDS sample loading buffer was added to beads, boiled at 95°C for 5 min, spun down and loaded on SDS-PAGE for western blot analysis. Western blotting was performed using standard methods and antibodies specific for 6x-HIS-Tag (Abcam), GFP (Santa Cruz), ITGB2 (R&D Systems), ITGB6 (R&D Systems), ITGA2 (R&D Systems), pMAPK (Cell Signaling), MAPK (Cell Signaling), pEGFR (Cell Signaling), EGFR (Cell Signaling), GAL3 (Santa Cruz), KRAS (Santa Cruz), pRAF1 (Cell signaling), RAF1 (Cell signaling), pERK (Cell signaling), ERK (Cell signaling), MYCN (Santa Cruz), TSG101 (Santa Cruz), CD63 (Santa Cruz), FN1 (Millipore), ACTA2 (Sigma), VIM (Cell Signaling), NKX2-1 (Santa Cruz), EZH2 (Abcam), H3K27me3 (Millipore), ASH-1 (Chemicon), TP53 (Cell signaling) and GAPDH (Sigma) were used. Immunoreactive proteins were visualized with the corresponding HRP-conjugated secondary antibodies (Santa Cruz) using the Super Signal West Femto detection solutions (ThermoFisher Scientific). Signals were detected and analyzed with Luminescent Image Analyzer (Las 4000, Fujifilm).

### KRAS activation assay

RAS Activation Assay Biochem Kit™ (BK008; Cytoskeleton, Inc) was used to assess KRAS activity following manufacturer’s instructions. Briefly, A549, NCI-H196 and NCI-H82 cell protein lysates were produced in cell lysis buffer containing 50 mM Tris–HCl pH 7.5, 10 mM MgCl2, 500 mM NaCl, 2% Igepal, 0–5% BSA, 20 mM Imidazole, 20 mM NaF, 0.5 mM Na3VO4, 40 μg/ml PMSF and protease inhibitor) for 10 min at 37 °C on rotator with 200 rpm. RAF-RBD beads (100 μg) were added to the reactions and incubated at 4 °C on a rotator for 1 h. The beads were washed once with 500 μl wash buffer (25 mM Tris pH 7.5, 30 mM MgCl2, 40 mM NaCl, 20 mM Imidazole, 20 mM NaF, 0.5 mM Na3VO4, 40 μg/ml PMSF and protease inhibitor). Precipitated proteins were analyzed by Western Blotting as described above.

### Proliferation Assay

NCI-196 cells were treated with Placebo, with binase or transfected with *siITGB2* and grown until 90–95% confluence in a 6-well plate. After 24 or 96h, cells were re-plated in 96-well plate in a density of 10^4^/well in a final volume of 100 μl and cultured in a humidified atmosphere at 37°C. 10 μM BrdU (Cell proliferation, colorimetric kit, Sigma #11647229001) were added and the cells were further incubated for additional 6 h, fixed and washed. Subsequently the immunoassay was done by measuring the absorbance of the samples in an ELISA reader at 370 nm (reference wavelength: 492 nm).

### Colony Assay

NCI-H196 cells transfected with *siCtrl* or *siITGB2* alone or in combination with binase treatment were plated in a 6-well culture plate at a density of 1000 or 5000 cells/well. The plate was swirled to ensure an even distribution of cells. The cells were grown in a 37 °C incubator with 5% CO2 for 3 to 5 days with media replacement every 2 days. At day 10, the media was removed and cells were washed twice with PBS. The colonies were analyzed using ImageJ software (https://imagej.nih.gov/ij/).

### Protein-interaction prediction

Prediction of protein-protein interaction was observed using the STRING online database ^69^ (Retrieval of Interacting Genes-Proteins–http://string.embl.de/) with a cut-off criterion of a combined score 0.9 (highest confidence) and including a maximum of 50 interactors on the 1^st^ shell and 20 interactors on the 2^nd^ shell. Network nodes represent proteins, while edges are the protein-protein associations. Small nodes represent protein of unknown 3D structure and large nodes proteins with some known or predicted 3D structures. Colored nodes represent query proteins and first shell of interactors and white nodes second shell of interactors. Interactions are depicted by color as follows: known interactions were obtained from curated databases (turquoise), or experimentally determined (purple); predicted interactions were defined by neighborhood (green), gene fusions (red) and gene co-occurrence (blue), textmining (light green), co-expression (black) and protein homology (violet). Top ITGA2 interactors were processed using the functional enrichment analyses Kyoto Encyclopedia of Genes and Genomes (KEGG). KEGG is the major public pathway-associated database, which identifies significantly enriched metabolic pathways or signal transduction pathways in target genes compared with the whole genome background.

### RNA sequencing and data analysis

RNA sequencing generated for this paper was sequenced as previously described ^67, 70^. Briefly, total RNA from A549 (Ctrl, *ITGB2* or *mutITGB2*) and NCI-H196 cells was isolated using the Trizol method. RNA was treated with DNase (DNase-Free DNase Set, Qiagen) and repurified using the RNeasy micro plus Kit (Qiagen). Total RNA and library integrity were verified on LabChip Gx Touch 24 (Perkin Elmer). Sequencing was performed on the NextSeq500 instrument (Illumina) using v2 chemistry with 1×75bp single end setup. Raw reads were visualized by FastQC to determine the quality of the sequencing. Trimming was performed using trimmomatic with the following parameters LEADING:3 TRAILING:3 SLIDINGWINDOW:4:15 HEADCROP:4, MINLEN:4. High quality reads were mapped using with STAR with reads corresponding to the transcript with default parameters. RNA-seq reads were mapped to human genome hg19. After mapping, Tag libraries were obtained with MakeTaglibrary from HOMER (default setting). Samples were quantified by using analyzeRepeats.pl with the parameters (hg19 -count genes –rpkm; reads per kilobase per millions mapped). Gene expression was quantified in reads per kilo base million (RPKM). Expression values of zero were set to the overall minimum value and all data were log2 transformed. Genes expressed (log2 transformed expression >0.2) were included in the analysis. The correlations of genes were measured using Pearson’s correlation. Overlapping genes in the cell lines RNAseq dataset were processed using the functional enrichment analyses KEGG, Gene Ontology and Reactome.

### RNA sequencing meta-analysis

RNAseq data of human lung adenocarcinoma (LUAD) (n=16) were downloaded from the TCGA data portal (https://tcga-data.nci.nih.gov/docs/publications/luad_2017/tcga.luad.rnaseq.20121025.csv.zip). The RNAseq data of SCLC and NSCLC cell lines was obtained from GEO under the accession number GSE30611 ^41^. 34 out of 675 samples were selected as NSCLC and 17 as SCLC for analysis based on the following sample annotations: “Organism part” is lung and “Diseases” is lung carcinoma, lung adenocarcinoma, lung anaplastic carcinoma, non-small cell lung carcinoma, squamous cell lung carcinoma, large cell lung carcinoma, lung mucoepidermoid carcinoma, lung papillary adenocarcinoma, lung adenosquamous carcinoma, bronchioloalveolar adenocarcinoma, or squamous cell carcinoma. For details on the original processing of the data, refer to the original paper ^37^. The transcriptome profile in NCI-H82 and NCI-196 was measured by the mean normalized expression of the genes in the A549 cell line. For analysis of RNA-seq data retrieved from the European Genome Archive ^71^, RNAseq was performed on cDNA libraries prepared from PolyA+ RNA extracted from SCLC tissues and cells. A library with an insert size of 250bp allowed to sequence 95bp paired-end reads without overlap. Raw reads were visualized by FastQC to determine the quality of the sequencing. Trimming was performed using trimmomatic with the following parameters LEADING:3 TRAILING:3 SLIDINGWINDOW:4:15 HEADCROP:4, MINLEN:4. High quality reads were mapped using with STAR with reads corresponding to the transcript with default parameters. RNA-seq reads were mapped to human genome hg19. After mapping, Tag libraries were obtained with MakeTaglibrary from HOMER (default setting). Samples were quantified by using analyzeRepeats.pl with the parameters (hg19 -count genes –rpkm; reads per kilobase per millions mapped). Gene expression was quantified in reads per kilo base million (RPKM).

### EVs purification, characterization and co-culture assays

For the collection of EVs, cells were cultured in media supplemented with 10% exosome-depleted FBS, in which EVs were depleted by overnight centrifugation at 100,000 g. Supernatants were then collected 72h later for EV purification. Cell culture supernatants were centrifuged at 500 g for 5 mins to pellet and discard cells, followed by 2,000 g for 30 min to remove cell debris and apoptotic bodies. A 1:1 volume of 2X PEG solution (16% w/v, polyethylene glycol, 1 M NaCl) was added. Samples were inverted to mix, then incubated overnight. Next day, medium/PEG mixture was centrifuged at 3,300 g for 1h. Crude vesicle pellets were resuspended in 1 ml of exosome-depleted 1X PBS and re-pelleted by centrifugation at 100,000 g for 70 minutes at 4°C (Beckman 45 Ti). Pellets at the bottom of the centrifugation tubes were resuspended in approximately 50ul of 1X PBS. Differential Light Scattering (DLS) was used to validate the EVs size range. WB was performed as described above to characterize the protein cargo of EVs using specific antibodies to detect EVs constitutive markers.

### Mass spectrometry: sample preparation, methods and data analysis

Extracellular vesicle samples were subjected to in gel digest as described ^72^. The resulting peptides were analyzed by liquid chromatography/tandem mass spectrometry (LC-MS2) utilizing in-house packed reverse phase column emitters (70 μm ID, 15 cm; ReproSil-Pur 120 C18-AQ, 1.9 μm, Dr. Maisch GmbH) and a buffer system comprising solvent A (5% acetonitrile, 0.1% formic acid) and solvent B (80% acetonitrile, 0.1% formic acid). The MaxQuant suite of algorithms (v.1.6.1.43) ^73–75^ was used for peptide/spectrum matching, protein group assembly as well as label free quantitation in the context of human Uniprot database (canonical and isoforms; downloaded on 2020/02/05; 210,349 entries). Relevant instrumentation parameters were extracted using MARMoSET ^76^ and are included in the supplementary material together with MaxQuant parametrization.

### Experiments with human PCLS

PCLS were prepared from tumor free lung explants from patients who underwent lung resection for cancer at KRH Hospital Siloah-Oststadt-Heidehaus or the Hanover Medical School (both Hanover, Germany). Tissue was processed immediately within one day of resection as described before ^77^. Briefly, human lung lobes were cannulated with a flexible catheter and the selected lung segments were inflated with warm (37°C) low melting agarose (1.5%) dissolved in Dulbecco’s Modified Eagle’s Medium Nutrient Mixture F-12 Ham (DMEM) supplemented with L-glutamine, 15 mM HEPES without phenol red, pH 7.2–7.4 (Sigma-Aldrich, Hamburg, Germany), 100 U/ml penicillin, and 100 µg/ml streptomycin (both from Biochrom, Berlin, Germany). After polymerization of the agarose solution on ice, tissue cores of a diameter of 8 mm were prepared using a sharp rotating metal tube. Subsequently, the cores were sliced into 300-350 µm thin slices in DMEM using a Krumdieck tissue slicer (Alabama Research and Development, Munford, AL). PCLS were washed 3× for 30 min in DMEM and used for experiments. Viability of the tissue was assessed by a LDH Cytotoxicity Detection Kit (Roche, Mannheim, Germany) according to manufacturer’s instruction.

For immunofluorescence staining in human PCLS, PCLS from Ctrl patients were fixed with acetone/methanol (Roth) 50:50 by volume for 20 min, blocked for 1 h with 5% bovine serum albumin (w/v, Sigma) in 1x PBS, pH 7.4. Cells were then incubated with primary antibody overnight at 4 °C. After incubation with a secondary antibody for 1h, nuclei were DAPI stained and PCLS were examined with a confocal microscope (Zeiss). Antibodies used were specific for ITGB2 (1:500 dilution, R&D Systems), and VIM (1:500 dilution, Cell Signaling). Alexa 488, Alexa555 or Alexa 594-tagged secondary antibodies (Invitrogen) were used. DAPI (Sigma Aldrich) used as nuclear dye.

### Statistical Analysis

The source data for all the plots presented in the article, including the values for statistical significance and the implemented statistical tests, are provided in Source Data S1. Further details of statistical analysis in different experiments are included in the Figures and Figure legends. Briefly, expression analysis of samples were analyzed by next generation sequencing in duplicates of one experiment. Three independent experiments of the mass spectrometry-based proteomic approach were performed. For the rest of the experiments presented here, samples were analyzed at least in triplicates and experiments were performed three times. Statistical analysis was performed using Excel Solver and Prism9. Data in bar plots are represented as mean ± standard error (mean ± s.e.m.). Two-tailed t-tests were used to determine the levels of difference between the groups and *P*-values for significance. *P*-values after two-tailed t-test, \**P* ≤ 0.05; \*\**P* < 0.01, and \*\*\**P* < 0.001.

### Data availability

The data that support this study are available from the corresponding author upon reasonable request. In addition, sequencing data of RNA have been deposited in NCBI’s Gene Expression Omnibus ^78^ and is accessible through SRA Sequence Read Archives NCBI with accession number PRJNA835424. The mass spectrometry-based interactome data have been deposited into the PRIDE archive and assigned to the project accession px-submission #576520. In addition, we retrieved and used a number of publicly available datasets to aid analysis of our data: Total RNA-seq in NSCLC and SCLC cell lines: European Genome Archive: EGAS00001000610 Total RNA-seq in SCLC patients and cell lines: European Genome Archive: EGAS00001002115, EGAS00001000299 The source data are provided with this paper.

## Acknowledgments

We thank Roswitha Bender for technical support; Julio Cordero, Stephanie Dobersch, Indrabahadur Singh, Ylia Salazar and Yanhan Xia for helpful discussions; Ernesto Guzmán-Díaz for FFPE lung tissue; Pourya Sarvari for human PCLS; Kerstin Richter and Alessandro Ianni for antibodies; Olga N Ilinskaya, Vera Ulyanova and Airat Kayumov for binase; Sylvie Fournel-Gigleux, Mohamed Ouzzine, Jean-Baptiste Vincourt, Sandrine Gulberti, Catherine Bui, Lydia Barré, Nick Ramalanjaona, Werner Seeger, Till Acker, Rajkumar Savai and Dulce Papy-Garcia for comments.

## Funding

Guillermo Barreto was funded by the “Centre National de la Recherche Scientifique” (CNRS, France), “Délégation Centre-Est” (CNRS-DR6) and the “Lorraine Université” (LU, France) through the initiative “Lorraine Université d’Excellence” (LUE) and the dispositive “Future Leader”, the Max-Planck-Society (MPG, Munich, Germany) and the “Deutsche Forschungsgemeinschaft” (DFG, Bonn, Germany) (BA 4036/4-1). Karla Rubio was funded by the “Consejo de Ciencia y Tecnología del Estado de Puebla” (CONCYTEP, Puebla, Mexico) through the initiative International Laboratory EPIGEN. Malgorzata Wygrecka acknowledges fundings from the DFG (WY119/1-3) and the German Center for Lung Research. Work in the lab of Thomas Braun is supported by the Deutsche Forschungsgemeinschaft, Excellence Cluster Cardio-Pulmonary Institute (CPI), Transregional Collaborative Research Center TRR81, TP A02, SFB1213 TP B02, TRR 267 TP A05 and the German Center for Cardiovascular Research. Karla Rubio received a doctoral fellowship from CONACyT-DAAD (PKZ91549687). Addi Romero received a doctoral fellowship from CONACyT-COCYT (CVU 510283).

## Author Contributions

KR, AJRO, PS, JG, SGü, AM and GB designed and performed the experiments; BB, PB, GD, MW, SGa and TB were involved in study design; GB, KR, AJRO, JG and SGü designed the study; GB, KR, AT, JG, SGü, PS and AJRO analyzed the data; GB, KR, JG, AT and AM wrote the manuscript. All authors discussed the results and commented on the manuscript.

## Conflict of Interest

The authors declare that they have no competing interests.

## Supplementary Information

**Figure S1:**
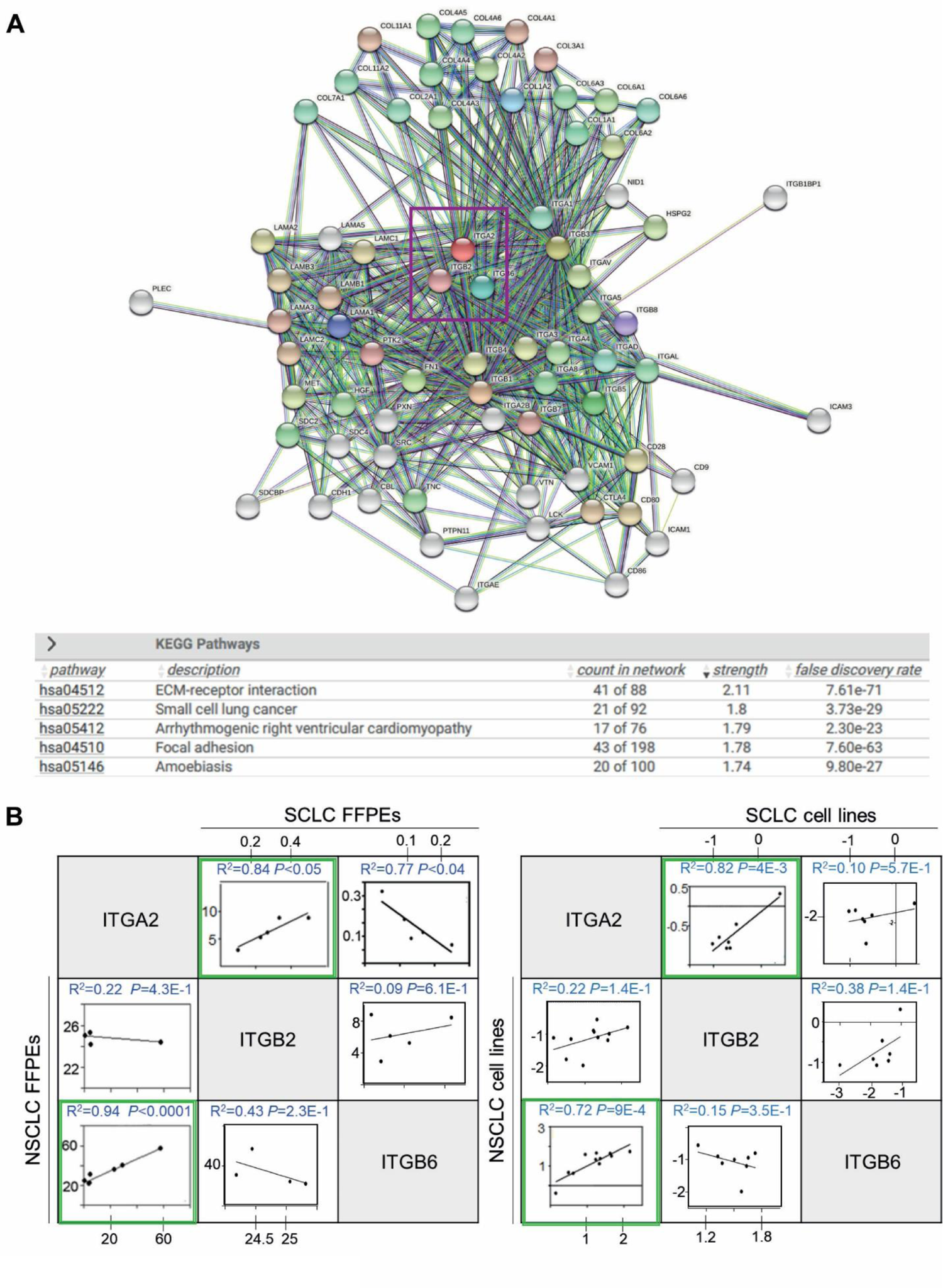
Human ITGA2 interactome is associated to SCLC. (**A**) In silico analysis of human ITGA2 interaction partners using the STRING 10.0 server. Pink box shows ITGA2, ITGB2, and ITGB6. Line colors indicate the known (turquoise), predicted (green), gene fusion (red), gene co-occurrence (blue) or experimental (purple) interactions. KEGG-based enrichment analysis of significant pathways including the ITGA2 interactome shows a significant enrichment with SCLC pathways. (**B**) Correlation analysis between *ITGA2*, *ITGB2* and *ITGB6* by linear regression of relative normalized expression in FFPE lung tissue sections from NSCLC and SCLC patients (left) and NSCLC and SCLC cells lines (right).

**Figure S2:**
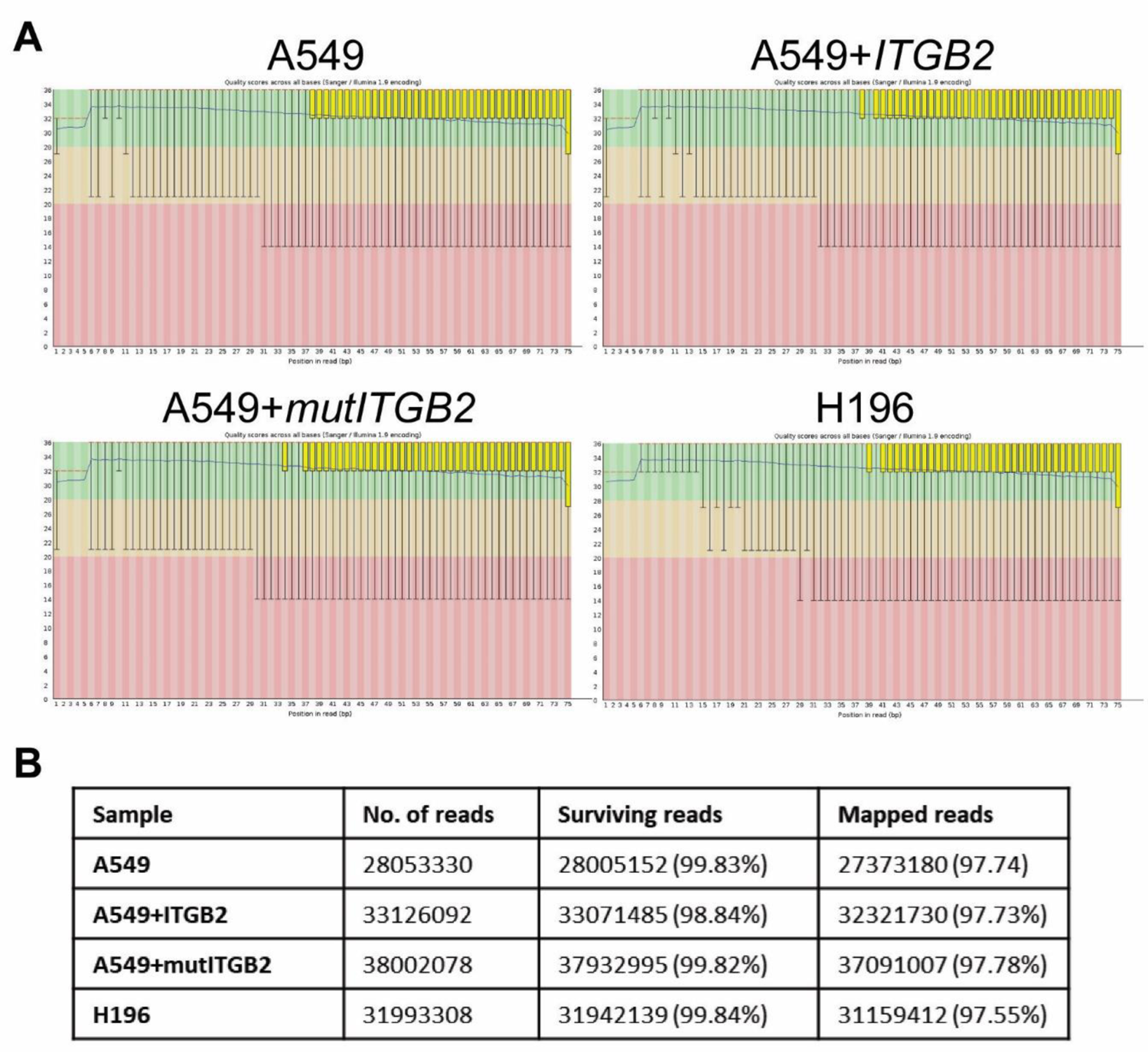
Transcriptomic analysis of SCLC and NSCLC ITGB2-overexpressing cells. (**A**) RNA-sequencing using SCLC (NCI-H196) and NSCLC (A549) cell lines without our with the overexpression of *ITGB2/mutITGB2* (accessible through the GEO BioProject ID PRJNA835424). Phred quality score distribution over all reads in each base. The score is divided into very good quality calls (green), calls of reasonable quality (orange), and calls of poor quality (red). (**B**) Description of the RNA-seq data sets supports the quality of the experiment.

**Figure S3:**
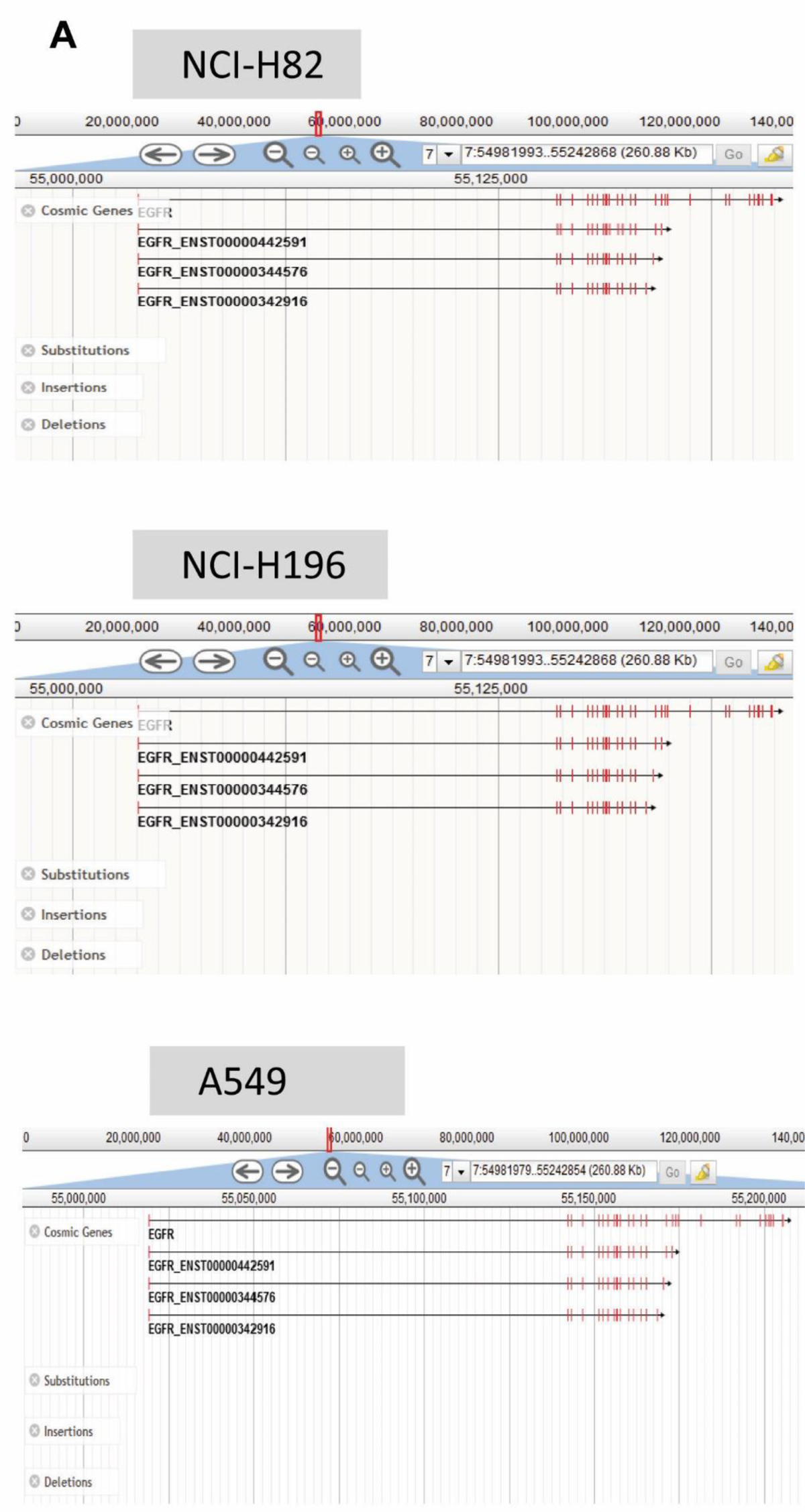
EGFR locus in NSCLC and SCLC cells. (**A**) Somatic mutations absence in the EGFR locus in NCI-H82, NCI-H196 and A549 cell lines from the Catalogue of Somatic Mutations in Cancer (COSMIC) is depicted. Mutation data were obtained from COSMIC v77 at the Welcome Trust Sanger Institute (Cambridge, UK). Only the frequency of somatic mutations (single nucleotides, or small insertions or deletions (indels)), but not larger deletions, amplifications or rearrangements, are considered.

**Figure S4:**
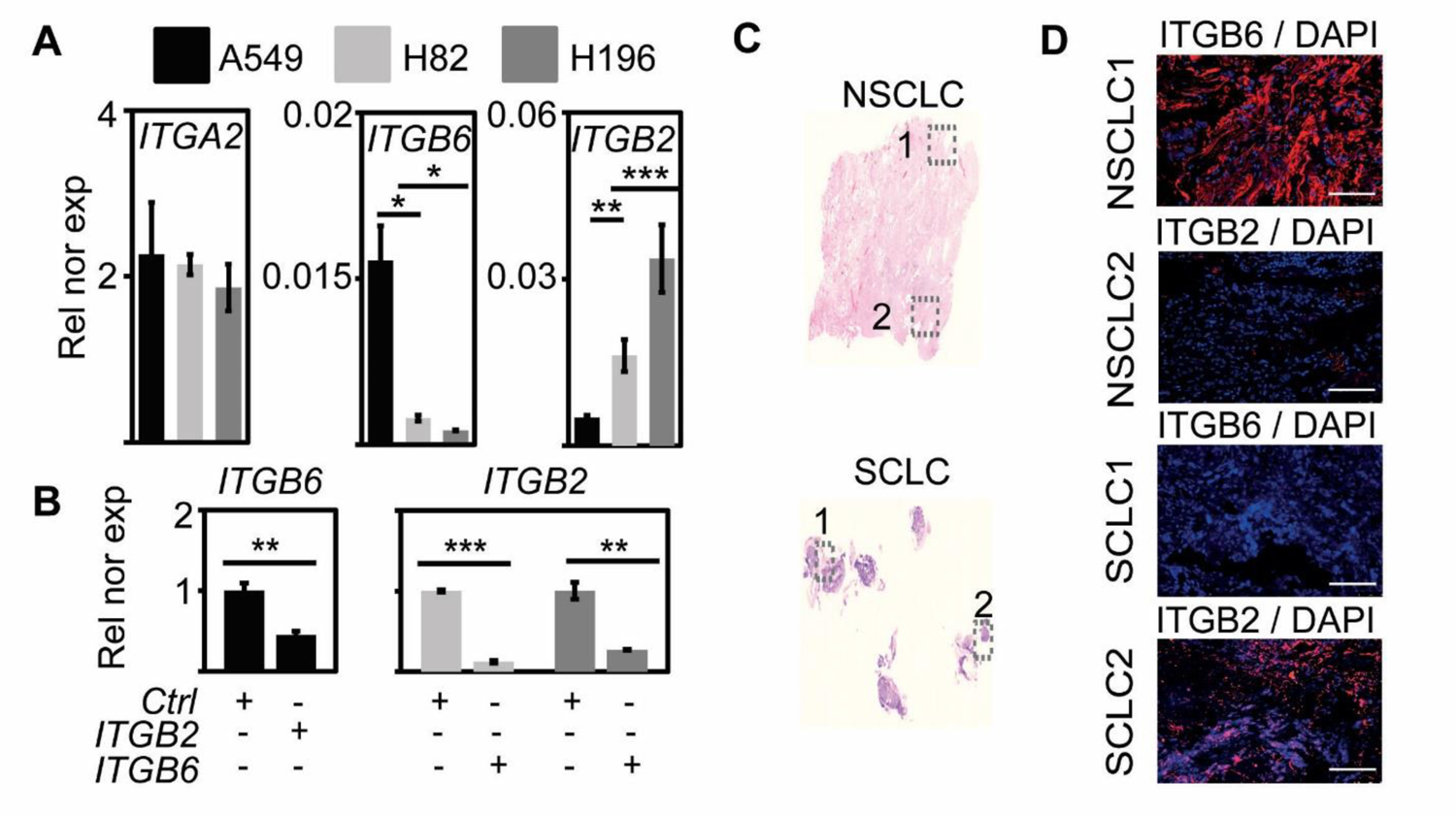
Mutual negative regulation of ITGB2 and ITGB6 levels in NSCLC and SCLC. (**A**) qRT-PCR-based expression analysis of indicated mRNA in A549, NCI-H82 and NCI-H196 cell lines. (**B**) qRT-PCR-based expression analysis of indicated mRNA in A549 cells (left panel) transfected with *ITGB2* or NCI-H196 cells (right panel) transfected with *ITGB6*. In the bar plots, data are shown as means ± s.e.m (*n*=3); asterisks, *P*-values after two-tailed t-test, \*\*\**P*˂0.001; \*\**P*˂0.01; \**P*˂0.05. (**C**) Hematoxylin and eosin staining in human lung tissue from NSCLC (top) and SCLC (bottom) patients. Squares are respectively shown in E at higher magnification. (**D**) Fluorescence microscopy after immunostaining using ITGB6 or ITGB2-specific antibodies in NSCLC and SCLC FFPE lung tissues (in C). DAPI, nuclear staining. Scale bar, 500 µm.

**Figure S5:**
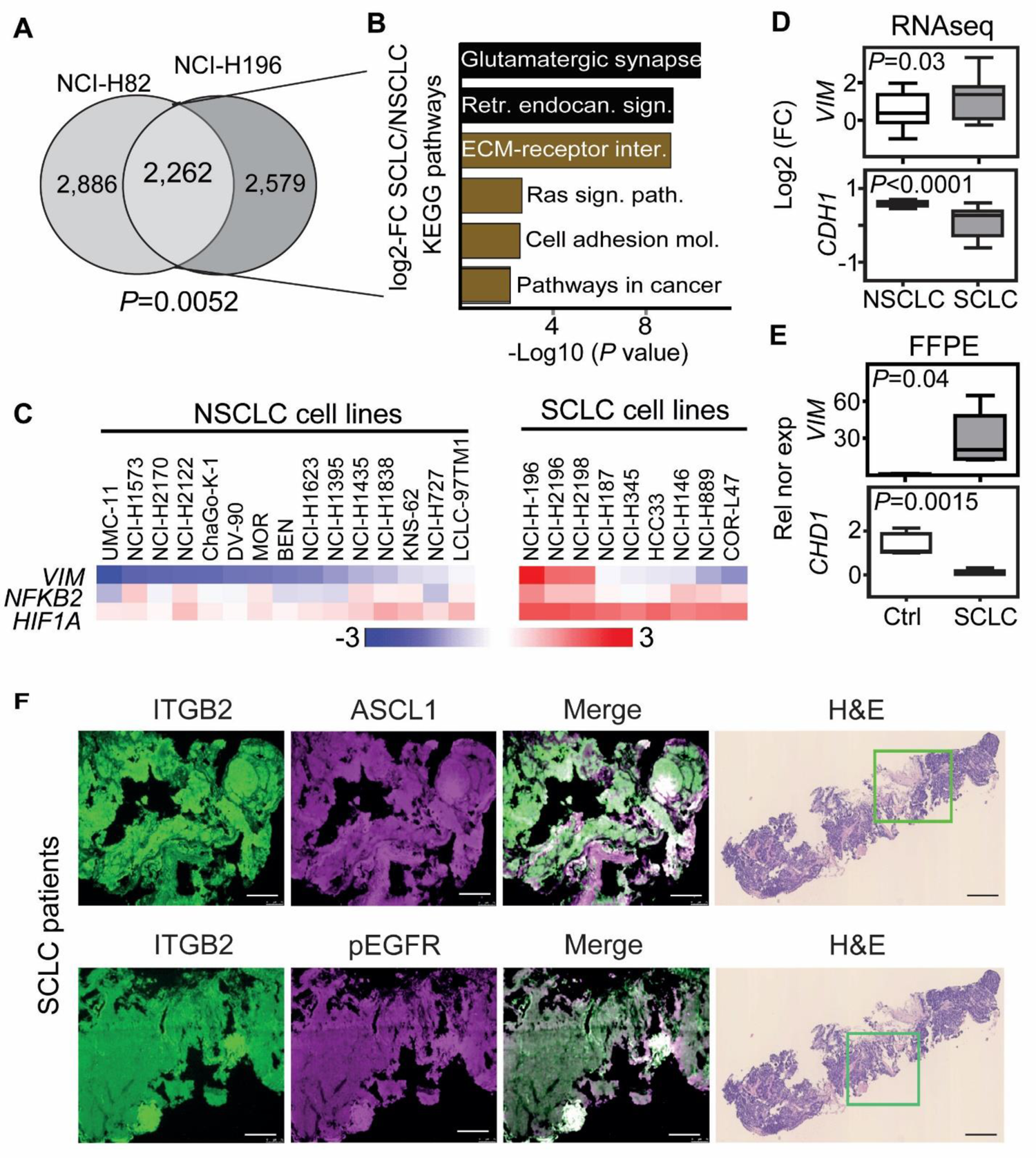
Active EGF signaling and pro-oncogenic pathways are predominant in ITGB2-expressing SCLC cell lines and lung tissue from SCLC patients. (**A**) Venn diagram displaying overlap of common, highly expressed genes in NCI-H82 and NCI-H196 cells when compared to A549 cells. *P*-value after Fisheŕs Exact test. (**B**) KEGG-based enrichment analysis of the common, highly expressed genes in NCI-H82 and NCI-H196 cells using DAVID bioinformatics tool and plotted by highest significance (-log10 of the modified Fisher exact *P*-value). Retr., retrograde; endocan., endocannabinoid; sign., signaling; path., pathway; mol. molecules. (**C**) Heatmaps of *VIM*, *NFKB2* and *HIF1A* expression in NSCLC and SCLC cancer cell lines. Hierarchical clustering was performed using Person’s correlation-based distance and average linkage. (**D**) Box plots of RNA-seq-based expression analysis of indicated transcripts in non-small cell lung cancer (NSCLC; *n*=33) and small cell lung cancer (SCLC; *n*=17) cell lines. Values are represented as log2 fold change (FC). (**E**) Box plots of qRT-PCR-based expression analysis of indicated transcripts using RNA isolated from FFPE lung tissue sections from Ctrl (*n*=4) and small cell-lung cancer (SCLC, *n*=5) patients. Rel nor exp, relative normalized expression to *GAPDH*. All box plots (D-E) indicate median (middle line), 25th, 75th percentile (box) and 5th and 95th percentile (whiskers); P-values after two-tailed t-test. Source data are provided as Source Data S1. (**F**) Left, fluorescence microscopy after immunostaining using ITGB2, ASH1 or pEGFR-specific antibodies in human lung tissue from SCLC patients. Right, hematoxylin and eosin staining (H&E) in human lung tissue from SCLC patients. Squares are respectively shown in left part of the panel at higher magnification. Scale bars, 150 µm (left) and 500 µm (right).

**Fig, S6:**
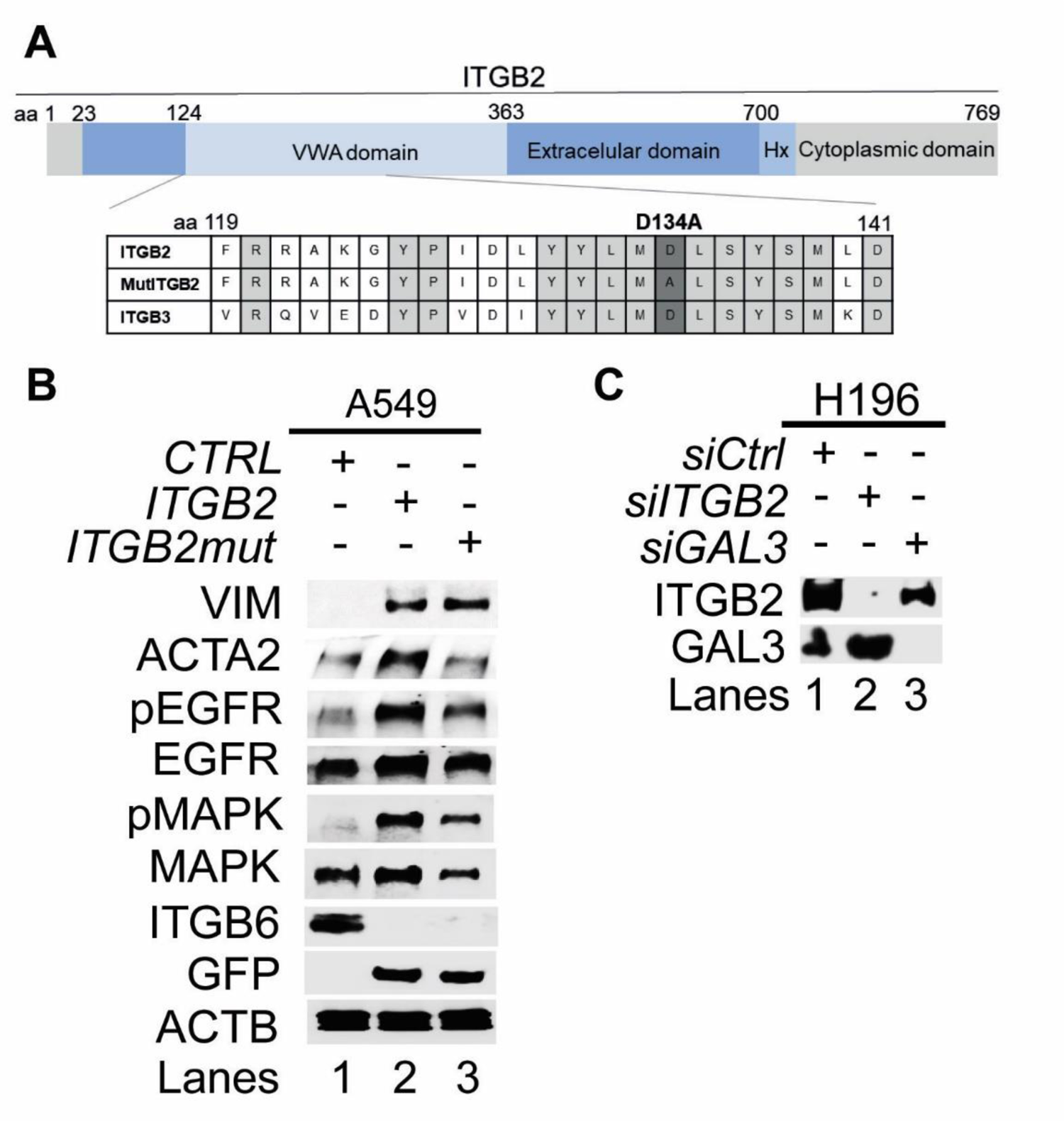
Non-canonical ITGB2 signaling activates EGFR. (**A**) Top, representation of ITGB2 functional domains. The numbers indicate amino acids (aa) positions. VWA, von Willebrand factor type A domain. Bottom, amino acid sequence alignment between ITGB2, mutITGB2 and ITGB3 in a part of the VWA domain highlighting the amino acid position 134, in which the point mutation D134A was incorporated to generate the ligand-binding-deficient ITGB2 mutant following a similar strategy as previously published for ITGB3 [PMID: 28860622]. (**B**) Total protein extracts of A549 cells transfected with *ITGB2* or *mutITGB2* were analyzed by WB using the indicated antibodies. (**C**) Total protein extracts from NCI-H196 cells that were transfected with Ctrl, ITGB2- or GAL3-specific small interfering RNAs (*siCtrl*, *siITGB2* or *siGAL3*) were analyzed by WB using the indicated antibodies.

**Figure S7:**
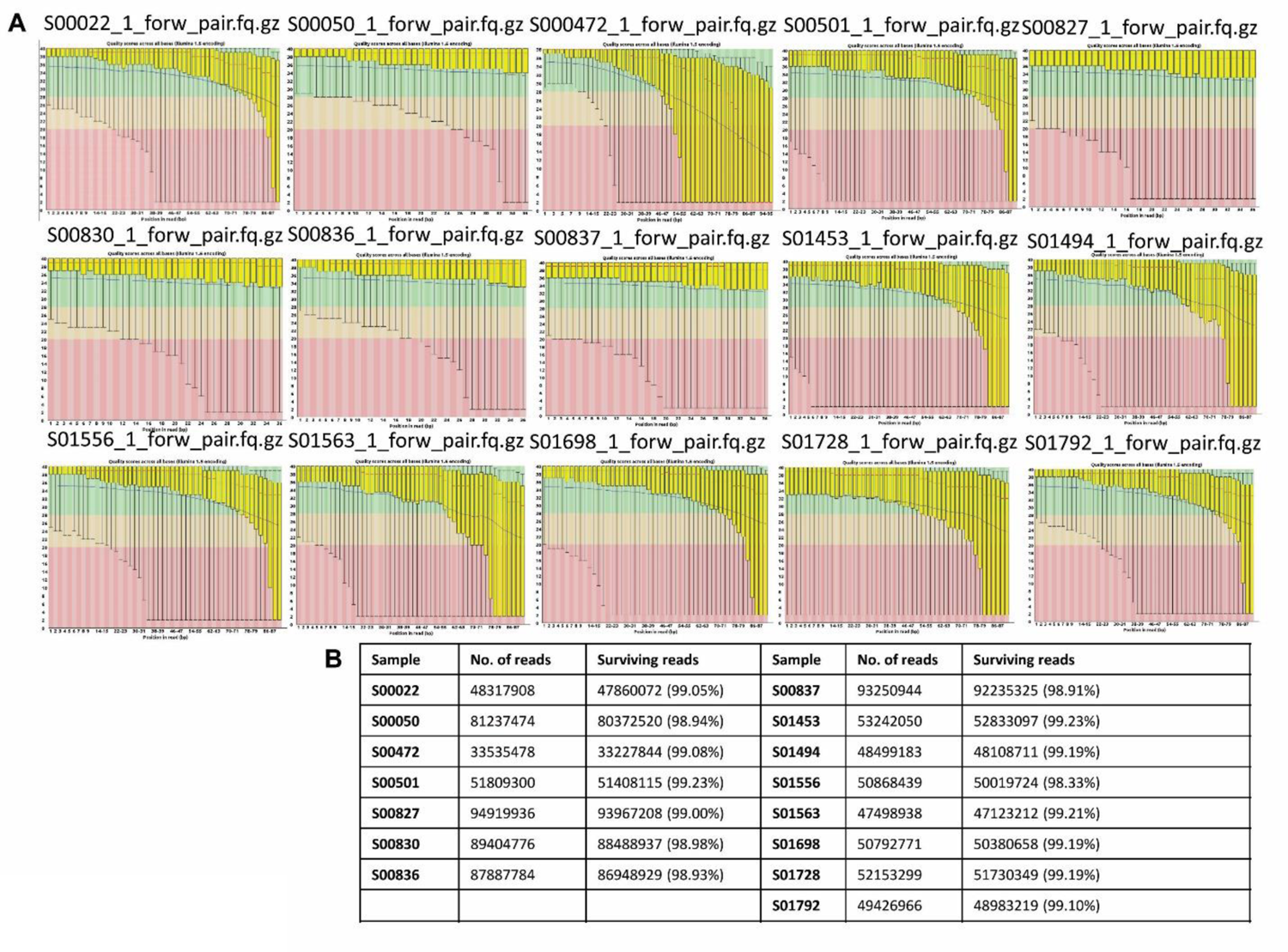
Transcriptomic analysis of SCLC patients tissue. (**A**), RNA-sequencing using lung tissue from SCLC patients (accessible through the European Genome Archive with accession number EGAS0000100299). Phred quality score distribution over all reads in each base. The score is divided into very good quality calls (green), calls of reasonable quality (orange), and calls of poor quality (red). (**B**), Description of the RNA-seq data sets supports the quality of the experiment.

**Figure S8:**
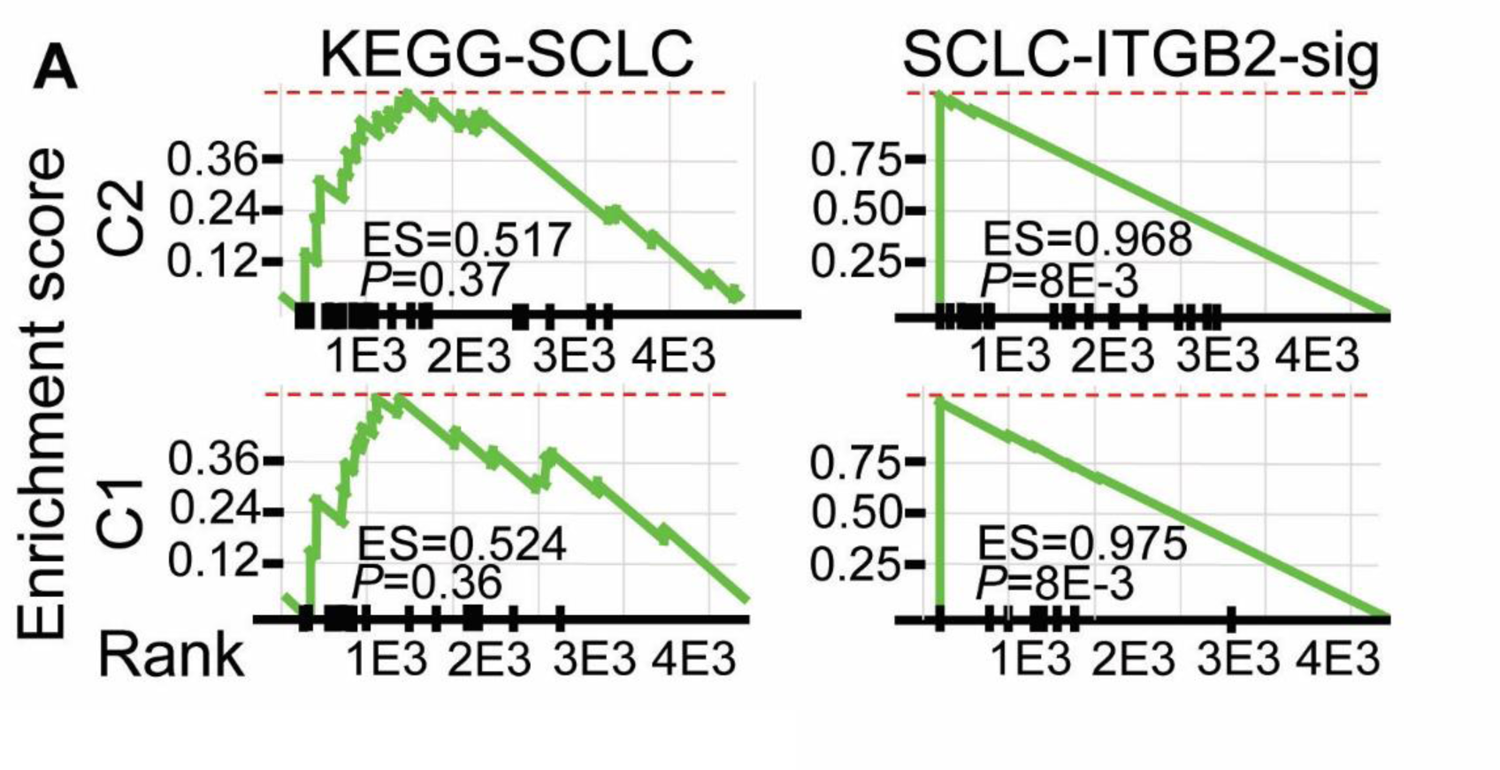
Confirmation of the SCLC-ITGB2 gene expression signature (SCLC-ITGB2-sig) using RNA-seq data from SCLC patients. Gene Set Enrichment Analysis (GSEA) using RNA-seq data from SCLC patients in cluster 1 (C1) and cluster 2 (C2) from Figure 2A comparing the conventional SCLC signature in KEGG (left) versus the SCLC-ITGB2-sig (right) identified in Figure 2E. ES, enrichment score; P-value after two-tailed t-test.

**Figure S9:**
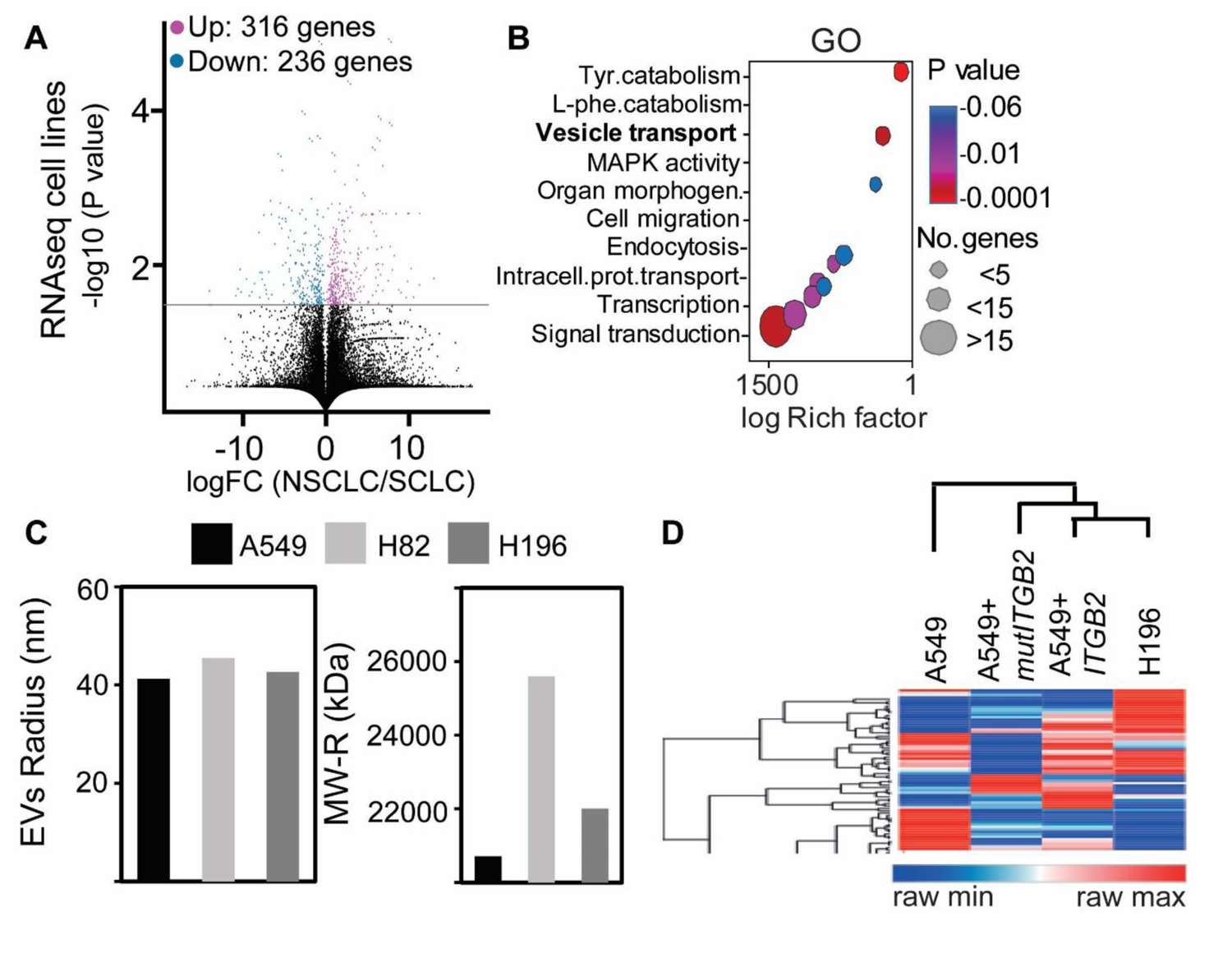
Extracellular vesicles containing ITGB2 are secreted from SCLC cells and ITGB2-transfected NSCLC cells. (**A**) Volcano plot representing the significance (-log10 *P*-values after two-tailed Welch’s t-Test) versus expression fold change (log2 expression ratios) between NSCLC and SCLC cells. Magenta dots show significantly upregulated transcripts, blue dots show significantly downregulated transcripts. (**B**) Gene Ontology-based enrichment analysis of up-regulated transcripts in SCLC using Webgestalt bioinformatics tool and plotted by highest significance (log Rich factor). Tyr., tyrosine; phe., phenylalanine; intracell., intracelular; prot., protein. (**C**) Radius (nm) and molecular weight (kDa) of extracellular vesicles (EVs) isolated from the culture medium of A549, NCI-H82 and NCI-H196 cells. Differential Light Scattering (DLS) was used to determine the EVs size. (**D**) Heatmap showing a hierarchical clustering from secreted EVs cargo proteins detected by Mass Spectrometry in supernatants of A549, A549+*ITGB2*, A549+*mutITGB2* and NCI-H196 cells. The complete proteomics data was submitted in the PRIDE repository with the accession number PX576520.

**Figure S10:**
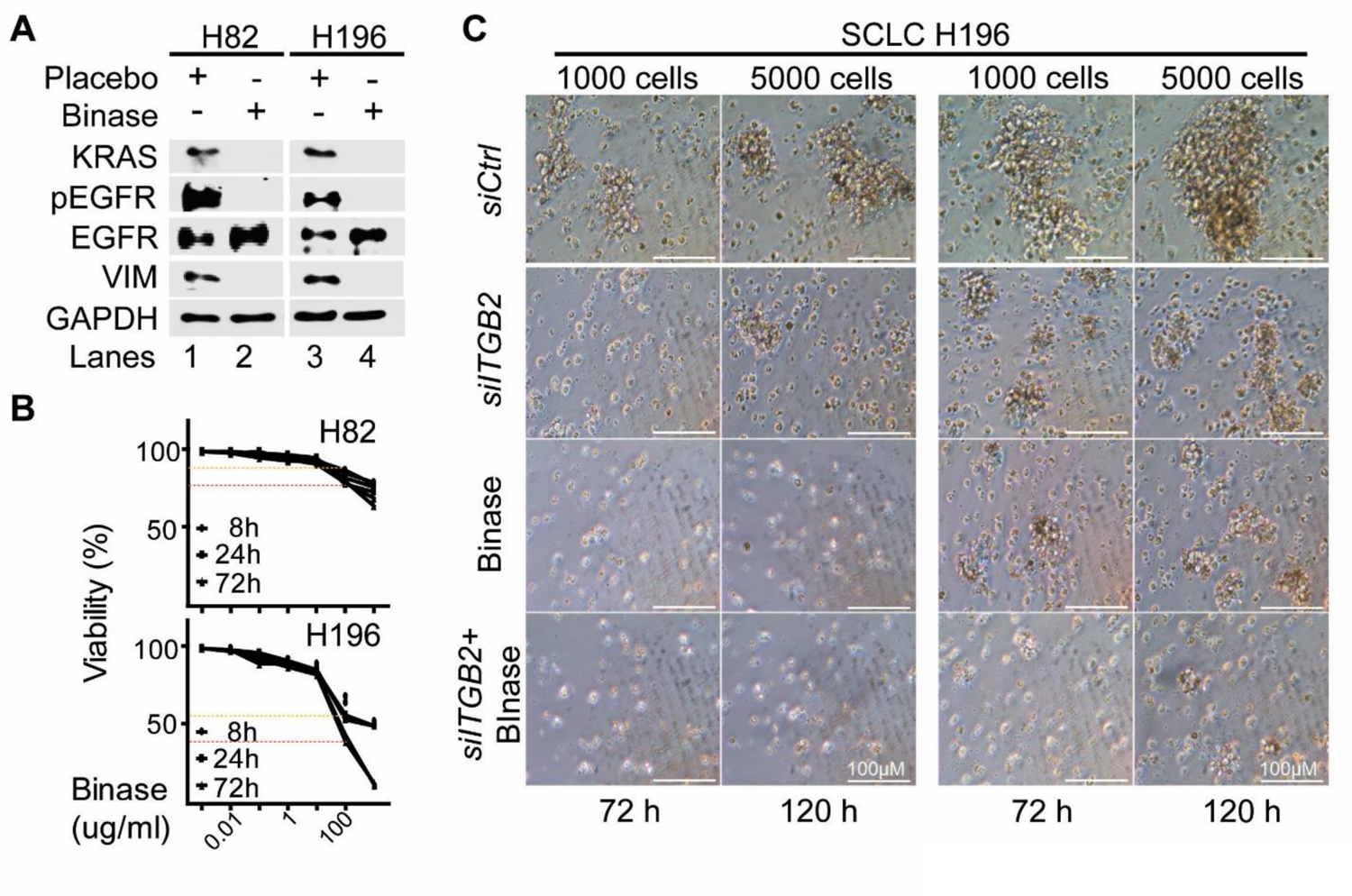
Binase counteracted the effects caused by EVs isolated from SCLC cell lines. (**A**) Total protein extracts of NCI-H82 and NCI-H196 cells treated with Placebo or with binase analyzed by WB using the indicated antibodies. (**B**) NCI-H82 and NCI-H196 cells were treated with increasing concentrations of binase for 8, 24 and 72h. Cell viability was determined using the BrdU incorporation fluorimetric assay. (**C**) Representative morphology of cell aggregates from NCI-H196 cells transfected with *siCtrl* or *siITGB2*, alone or in combination with binase treatment. The images represent the change of morphology of cell aggregates after aggregate formation for 72h and 120h with initial 1000 or 5000 cells.

### Data S1. (separate files)

Source Data S1 - This is an Excel file that contains the data for all the plots presented in the article, including the values for statistical significance and the implemented statistical tests.

Tables S1 – This is an Excel file that contains the 50 interaction partners of human ITGA2 with high confidence (combined score ≥ 0.9; 2 nodes), including ITGB2 (combined score=0.96) and ITGB6 (combined score=0.97), that we identified using the STRING database.

Table S2 – This is an Excel file that contains the clinical and pathological characteristics of SCLC and Ctrl patients, from which we obtained the formalin-fixed paraffin embedded (FFPE) human lung tissues.

Table S3 - This is an Excel file that contains the clinical and pathological characteristics of LUAD and Ctrl patients, from which we retrieved RNA-seq data from The Cancer Genome Atlas (TCGA).

Table S4 – This is an Excel file that contains the IDs of the transcripts that we suggest as SCLC- ITGB2 gene expression signature (SCLC-ITGB2-sig).

Table S5 – This is an Excel file that contains the official symbols of the 189 proteins that we have identified by mass spectrometry analysis of the protein cargo of the isolated EVs from NCI- H196 cells transfected with control plasmid (*Ctrl*) and from A549 cells transfected either with *ITGB2* or *mutITGB2* (Figure 5D and Figure S9D)

## Notes

### Competing Interest Statement

The authors have declared no competing interest.

### Summary of Updates

Corresponding authorship (1 online, 2 in PDF)

## References

1 Mehta, A., Dobersch, S., Romero-Olmedo, A. J. & Barreto, G. Epigenetics in lung cancer diagnosis and therapy. Cancer metastasis reviews, doi:10.1007/s10555-015-9563-3 (2015).

2 Blandin Knight, S. et al. Progress and prospects of early detection in lung cancer. Open Biol 7, doi:10.1098/rsob.170070 (2017).

3 Herbst, R. S., Heymach, J. V. & Lippman, S. M. Lung cancer. The New England journal of medicine 359, 1367–1380, doi:10.1056/NEJMra0802714 (2008).

4 Sabari, J. K., Lok, B. H., Laird, J. H., Poirier, J. T. & Rudin, C. M. Unravelling the biology of SCLC: implications for therapy. Nat Rev Clin Oncol 14, 549–561, doi:10.1038/nrclinonc.2017.71 (2017).

5 Tartarone, A. et al. Progress and challenges in the treatment of small cell lung cancer. Medical oncology 34, 110, doi:10.1007/s12032-017-0966-6 (2017).

6 Thery, C., Amigorena, S., Raposo, G. & Clayton, A. Isolation and characterization of exosomes from cell culture supernatants and biological fluids. Current protocols in cell biology / editorial board, Juan S. Bonifacino … [et al.] Chapter 3, Unit 3 22, doi:10.1002/0471143030.cb0322s30 (2006).

7 Brennan, K. et al. A comparison of methods for the isolation and separation of extracellular vesicles from protein and lipid particles in human serum. Scientific reports 10, 1039, doi:10.1038/s41598-020-57497-7 (2020).

8 Thery, C., Ostrowski, M. & Segura, E. Membrane vesicles as conveyors of immune responses. Nat Rev Immunol 9, 581–593, doi:10.1038/nri2567 (2009).

9 Hristov, M., Erl, W., Linder, S. & Weber, P. C. Apoptotic bodies from endothelial cells enhance the number and initiate the differentiation of human endothelial progenitor cells in vitro. Blood 104, 2761–2766, doi:10.1182/blood-2003-10-3614 (2004).

10 Yanez-Mo, M. et al. Biological properties of extracellular vesicles and their physiological functions. J Extracell Vesicles 4, 27066, doi:10.3402/jev.v4.27066 (2015).

11 Booth, A. M. et al. Exosomes and HIV Gag bud from endosome-like domains of the T cell plasma membrane. The Journal of cell biology 172, 923–935, doi:10.1083/jcb.200508014 (2006).

12 Mastoridis, S. et al. Multiparametric Analysis of Circulating Exosomes and Other Small Extracellular Vesicles by Advanced Imaging Flow Cytometry. Front Immunol 9, 1583, doi:10.3389/fimmu.2018.01583 (2018).

13 van Niel, G. et al. Intestinal epithelial cells secrete exosome-like vesicles. Gastroenterology 121, 337–349, doi:10.1053/gast.2001.26263 (2001).

14 Borges, F. T. et al. TGF-beta1-containing exosomes from injured epithelial cells activate fibroblasts to initiate tissue regenerative responses and fibrosis. Journal of the American Society of Nephrology: JASN 24, 385–392, doi:10.1681/ASN.2012101031 (2013).

15 Panigrahi, G. K. et al. Hypoxia-induced exosome secretion promotes survival of African-American and Caucasian prostate cancer cells. Scientific reports 8, 3853, doi:10.1038/s41598-018-22068-4 (2018).

16 Boussadia, Z. et al. Acidic microenvironment plays a key role in human melanoma progression through a sustained exosome mediated transfer of clinically relevant metastatic molecules. J Exp Clin Cancer Res 37, 245, doi:10.1186/s13046-018-0915-z (2018).

17 Hoshino, A. et al. Extracellular Vesicle and Particle Biomarkers Define Multiple Human Cancers. Cell 182, 1044–1061 e1018, doi:10.1016/j.cell.2020.07.009 (2020).

18 Hurwitz, S. N. & Meckes, D. G., Jr. Extracellular Vesicle Integrins Distinguish Unique Cancers. Proteomes 7, doi:10.3390/proteomes7020014 (2019).

19 Chalmin, F. et al. Membrane-associated Hsp72 from tumor-derived exosomes mediates STAT3-dependent immunosuppressive function of mouse and human myeloid-derived suppressor cells. The Journal of clinical investigation 120, 457–471, doi:10.1172/JCI40483 (2010).

20 Hsu, Y. L. et al. Hypoxic lung cancer-secreted exosomal miR-23a increased angiogenesis and vascular permeability by targeting prolyl hydroxylase and tight junction protein ZO-1. Oncogene 36, 4929–4942, doi:10.1038/onc.2017.105 (2017).

21 Tadokoro, H., Umezu, T., Ohyashiki, K., Hirano, T. & Ohyashiki, J. H. Exosomes derived from hypoxic leukemia cells enhance tube formation in endothelial cells. The Journal of biological chemistry 288, 34343–34351, doi:10.1074/jbc.M113.480822 (2013).

22 Umezu, T. et al. Exosomal miR-135b shed from hypoxic multiple myeloma cells enhances angiogenesis by targeting factor-inhibiting HIF-1. Blood 124, 3748–3757, doi:10.1182/blood-2014-05-576116 (2014).

23 Pelissier Vatter, F. A. et al. Extracellular vesicle- and particle-mediated communication shapes innate and adaptive immune responses. The Journal of experimental medicine 218, doi:10.1084/jem.20202579 (2021).

24 Liu, T. C., Jin, X., Wang, Y. & Wang, K. Role of epidermal growth factor receptor in lung cancer and targeted therapies. Am J Cancer Res 7, 187–202 (2017).

25 Takeuchi, K. & Ito, F. EGF receptor in relation to tumor development: molecular basis of responsiveness of cancer cells to EGFR-targeting tyrosine kinase inhibitors. The FEBS journal 277, 316–326, doi:10.1111/j.1742-4658.2009.07450.x (2010).

26 Martinelli, E., Morgillo, F., Troiani, T. & Ciardiello, F. Cancer resistance to therapies against the EGFR-RAS-RAF pathway: The role of MEK. Cancer Treat Rev 53, 61–69, doi:10.1016/j.ctrv.2016.12.001 (2017).

27 Campbell, I. D. & Humphries, M. J. Integrin structure, activation, and interactions. Cold Spring Harbor perspectives in biology 3, doi:10.1101/cshperspect.a004994 (2011).

28 Hynes, R. O. Integrins: bidirectional, allosteric signaling machines. Cell 110, 673–687 (2002).

29 Munger, J. S. & Sheppard, D. Cross talk among TGF-beta signaling pathways, integrins, and the extracellular matrix. Cold Spring Harbor perspectives in biology 3, a005017, doi:10.1101/cshperspect.a005017 (2011).

30 Mori, S. et al. Direct binding of integrin alphavbeta3 to FGF1 plays a role in FGF1 signaling. The Journal of biological chemistry 283, 18066–18075, doi:10.1074/jbc.M801213200 (2008).

31 Guo, W. & Giancotti, F. G. Integrin signalling during tumour progression. Nature reviews. Molecular cell biology 5, 816–826, doi:10.1038/nrm1490 (2004).

32 Carpenter, B. L. et al. Integrin alpha6beta4 Promotes Autocrine Epidermal Growth Factor Receptor (EGFR) Signaling to Stimulate Migration and Invasion toward Hepatocyte Growth Factor (HGF). The Journal of biological chemistry 290, 27228–27238, doi:10.1074/jbc.M115.686873 (2015).

33 Seguin, L. et al. An integrin beta(3)-KRAS-RalB complex drives tumour stemness and resistance to EGFR inhibition. Nature cell biology 16, 457–468, doi:10.1038/ncb2953 (2014).

34 Ricono, J. M. et al. Specific cross-talk between epidermal growth factor receptor and integrin alphavbeta5 promotes carcinoma cell invasion and metastasis. Cancer research 69, 1383–1391, doi:10.1158/0008-5472.CAN-08-3612 (2009).

35 Mukhametshina, R. T. et al. Quantitative proteome analysis of alveolar type-II cells reveals a connection of integrin receptor subunits beta 2/6 and WNT signaling. Journal of proteome research 12, 5598–5608, doi:10.1021/pr400573k (2013).

36 Xu, X. et al. Evidence for type II cells as cells of origin of K-Ras-induced distal lung adenocarcinoma. Proceedings of the National Academy of Sciences of the United States of America 109, 4910–4915, doi:10.1073/pnas.1112499109 (2012).

37 Klijn, C. et al. A comprehensive transcriptional portrait of human cancer cell lines. Nature biotechnology 33, 306–312, doi:10.1038/nbt.3080 (2015).

38 Lim, S. Y. et al. LSD1 modulates the non-canonical integrin beta3 signaling pathway in non-small cell lung carcinoma cells. Scientific reports 7, 10292, doi:10.1038/s41598-017-09554-x (2017).

39 George, J. et al. Comprehensive genomic profiles of small cell lung cancer. Nature 524, 47–53, doi:10.1038/nature14664 (2015).

40 Jassal, B. et al. The reactome pathway knowledgebase. Nucleic acids research 48, D498–D503, doi:10.1093/nar/gkz1031 (2020).

41 Park, Y., Lim, S., Nam, J. W. & Kim, S. Measuring intratumor heterogeneity by network entropy using RNA-seq data. Scientific reports 6, 37767, doi:10.1038/srep37767 (2016).

42 Lanczky, A. & Gyorffy, B. Web-Based Survival Analysis Tool Tailored for Medical Research (KMplot): Development and Implementation. J Med Internet Res 23, e27633, doi:10.2196/27633 (2021).

43 Stephen, A. G., Esposito, D., Bagni, R. K. & McCormick, F. Dragging ras back in the ring. Cancer cell 25, 272–281, doi:10.1016/j.ccr.2014.02.017 (2014).

44 Ilinskaya, O. N. et al. Direct inhibition of oncogenic KRAS by Bacillus pumilus ribonuclease (binase). Biochimica et biophysica acta 1863, 1559–1567, doi:10.1016/j.bbamcr.2016.04.005 (2016).

45 Kilinc, S. et al. Oncogene-regulated release of extracellular vesicles. Developmental cell 56, 1989–2006 e1986, doi:10.1016/j.devcel.2021.05.014 (2021).

46 Thery, C. et al. Minimal information for studies of extracellular vesicles 2018 (MISEV2018): a position statement of the International Society for Extracellular Vesicles and update of the MISEV2014 guidelines. J Extracell Vesicles 7, 1535750, doi:10.1080/20013078.2018.1535750 (2018).

47 Piper, R. C. & Katzmann, D. J. Biogenesis and function of multivesicular bodies. Annual review of cell and developmental biology 23, 519–547, doi:10.1146/annurev.cellbio.23.090506.123319 (2007).

48 Poirier, J. T. et al. DNA methylation in small cell lung cancer defines distinct disease subtypes and correlates with high expression of EZH2. Oncogene 34, 5869–5878, doi:10.1038/onc.2015.38 (2015).

49 Mollaoglu, G. et al. MYC Drives Progression of Small Cell Lung Cancer to a Variant Neuroendocrine Subtype with Vulnerability to Aurora Kinase Inhibition. Cancer cell 31, 270–285, doi:10.1016/j.ccell.2016.12.005 (2017).

50 Azuma, K. et al. Phase II study of erlotinib plus tivantinib (ARQ 197) in patients with locally advanced or metastatic EGFR mutation-positive non-small-cell lung cancer just after progression on EGFR-TKI, gefitinib or erlotinib. ESMO Open 1, e000063, doi:10.1136/esmoopen-2016-000063 (2016).

51 van der Wekken, A. J. et al. Overall survival in EGFR mutated non-small-cell lung cancer patients treated with afatinib after EGFR TKI and resistant mechanisms upon disease progression. PloS one 12, e0182885, doi:10.1371/journal.pone.0182885 (2017).

52 Suda, K., Onozato, R., Yatabe, Y. & Mitsudomi, T. EGFR T790M mutation: a double role in lung cancer cell survival? J Thorac Oncol 4, 1–4, doi:10.1097/JTO.0b013e3181913c9f (2009).

53 Kobayashi, S. et al. EGFR mutation and resistance of non-small-cell lung cancer to gefitinib. The New England journal of medicine 352, 786–792, doi:10.1056/NEJMoa044238 (2005).

54 Sequist, L. V. et al. Genotypic and histological evolution of lung cancers acquiring resistance to EGFR inhibitors. Science translational medicine 3, 75ra26, doi:10.1126/scitranslmed.3002003 (2011).

55 Haas, T. L. et al. Integrin alpha7 Is a Functional Marker and Potential Therapeutic Target in Glioblastoma. Cell stem cell 21, 35–50 e39, doi:10.1016/j.stem.2017.04.009 (2017).

56 Naci, D. et al. alpha2beta1 integrin promotes chemoresistance against doxorubicin in cancer cells through extracellular signal-regulated kinase (ERK). The Journal of biological chemistry 287, 17065–17076, doi:10.1074/jbc.M112.349365 (2012).

57 Kim, Y. J., Jung, K., Baek, D. S., Hong, S. S. & Kim, Y. S. Co-targeting of EGF receptor and neuropilin-1 overcomes cetuximab resistance in pancreatic ductal adenocarcinoma with integrin beta1-driven Src-Akt bypass signaling. Oncogene 36, 2543–2552, doi:10.1038/onc.2016.407 (2017).

58 Lishko, V. K., Yakubenko, V. P. & Ugarova, T. P. The interplay between integrins alphaMbeta2 and alpha5beta1 during cell migration to fibronectin. Exp Cell Res 283, 116–126 (2003).

59 Rainero, E. et al. Diacylglycerol kinase alpha controls RCP-dependent integrin trafficking to promote invasive migration. The Journal of cell biology 196, 277–295, doi:10.1083/jcb.201109112 (2012).

60 Hang, Q. et al. N-Glycosylation of integrin alpha5 acts as a switch for EGFR-mediated complex formation of integrin alpha5beta1 to alpha6beta4. Scientific reports 6, 33507, doi:10.1038/srep33507 (2016).

61 Wang, H., Jin, H. & Rapraeger, A. C. Syndecan-1 and Syndecan-4 Capture Epidermal Growth Factor Receptor Family Members and the alpha3beta1 Integrin Via Binding Sites in Their Ectodomains: NOVEL SYNSTATINS PREVENT KINASE CAPTURE AND INHIBIT alpha6beta4-INTEGRIN-DEPENDENT EPITHELIAL CELL MOTILITY. The Journal of biological chemistry 290, 26103–26113, doi:10.1074/jbc.M115.679084 (2015).

62 Shah, L. et al. Expression of syndecan-1 and expression of epidermal growth factor receptor are associated with survival in patients with nonsmall cell lung carcinoma. Cancer 101, 1632–1638, doi:10.1002/cncr.20542 (2004).

63 Hang, Q. et al. Integrin alpha5 Suppresses the Phosphorylation of Epidermal Growth Factor Receptor and Its Cellular Signaling of Cell Proliferation via N-Glycosylation. The Journal of biological chemistry 290, 29345–29360, doi:10.1074/jbc.M115.682229 (2015).

64 Isaji, T. et al. N-glycosylation of the beta-propeller domain of the integrin alpha5 subunit is essential for alpha5beta1 heterodimerization, expression on the cell surface, and its biological function. The Journal of biological chemistry 281, 33258–33267, doi:10.1074/jbc.M607771200 (2006).

65 Kariya, Y. & Gu, J. N-glycosylation of ss4 integrin controls the adhesion and motility of keratinocytes. PloS one 6, e27084, doi:10.1371/journal.pone.0027084 (2011).

66 Mehta, A. et al. Validation of Tuba1a as Appropriate Internal Control for Normalization of Gene Expression Analysis during Mouse Lung Development. International journal of molecular sciences 16, 4492–4511, doi:10.3390/ijms16034492 (2015).

67 Rubio, K. et al. Inactivation of nuclear histone deacetylases by EP300 disrupts the MiCEE complex in idiopathic pulmonary fibrosis. Nature communications 10, 2229, doi:10.1038/s41467-019-10066-7 (2019).

68 Singh, I. et al. High mobility group protein-mediated transcription requires DNA damage marker gamma-H2AX. Cell research 25, 837–850, doi:10.1038/cr.2015.67 (2015).

69 Szklarczyk, D. et al. The STRING database in 2017: quality-controlled protein-protein association networks, made broadly accessible. Nucleic acids research 45, D362–D368, doi:10.1093/nar/gkw937 (2017).

70 Singh, I. et al. MiCEE is a ncRNA-protein complex that mediates epigenetic silencing and nucleolar organization. Nature genetics 50, 990–1001, doi:10.1038/s41588-018-0139-3 (2018).

71 Peifer, M. et al. Integrative genome analyses identify key somatic driver mutations of small-cell lung cancer. Nature genetics 44, 1104–1110, doi:10.1038/ng.2396 (2012).

72 Schutt, C. et al. Linc-MYH configures INO80 to regulate muscle stem cell numbers and skeletal muscle hypertrophy. The EMBO journal 39, e105098, doi:10.15252/embj.2020105098 (2020).

73 Cox, J. & Mann, M. MaxQuant enables high peptide identification rates, individualized p.p.b.-range mass accuracies and proteome-wide protein quantification. Nature biotechnology 26, 1367–1372, doi:10.1038/nbt.1511 (2008).

74 Cox, J. et al. Andromeda: a peptide search engine integrated into the MaxQuant environment. Journal of proteome research 10, 1794–1805, doi:10.1021/pr101065j (2011).

75 Cox, J. et al. Accurate proteome-wide label-free quantification by delayed normalization and maximal peptide ratio extraction, termed MaxLFQ. Molecular & cellular proteomics: MCP 13, 2513–2526, doi:10.1074/mcp.M113.031591 (2014).

76 Kiweler, M., Looso, M. & Graumann, J. MARMoSET - Extracting Publication-ready Mass Spectrometry Metadata from RAW Files. Molecular & cellular proteomics: MCP 18, 1700–1702, doi:10.1074/mcp.TIR119.001505 (2019).

77 Dobersch, S. et al. Positioning of nucleosomes containing gamma-H2AX precedes active DNA demethylation and transcription initiation. Nature communications 12, 1072, doi:10.1038/s41467-021-21227-y (2021).

78 Edgar, R., Domrachev, M. & Lash, A. E. Gene Expression Omnibus: NCBI gene expression and hybridization array data repository. Nucleic acids research 30, 207–210 (2002).

